# An integrative analysis of genomic and exposomic data for complex traits and phenotypic prediction

**DOI:** 10.1101/2020.11.09.373704

**Authors:** Xuan Zhou, S. Hong Lee

**Affiliations:** Australian Centre for Precision Health, University of South Australia, Adelaide, South Australia, 5000, Australia; UniSA Allied Health and Human Performance, University of South Australia, Adelaide, South Australia, 5000, Australia; South Australian Health and Medical Research Institute, Adelaide, South Australia, 5000, Australia

## Abstract

Complementary to the genome, the concept of exposome has been proposed to capture the totality of human environmental exposures. While there has been some recent progress on the construction of the exposome, few tools exist that can integrate the genome and exposome for complex trait analyses. Here we propose a linear mixed model approach to bridge this gap, which jointly models the random effects of the two omics layers on phenotypes of complex traits. We illustrate our approach using traits from the UK Biobank (e.g., BMI & height for N ∼ 35,000) with a small fraction of the exposome that comprises 28 lifestyle factors. The joint model of the genome and exposome explains substantially more phenotypic variance and significantly improves phenotypic prediction accuracy, compared to the model based on the genome alone. The additional phenotypic variance captured by the exposome includes its additive effects as well as non-additive effects such as genome-exposome (gxe) and exposome-exposome (exe) interactions. For example, 19% of variation in BMI is explained by additive effects of the genome, while additional 7.2% by additive effects of the exposome, 1.9% by exe interactions and 4.5% by gxe interactions. Correspondingly, the prediction accuracy for BMI, computed using Pearson’s correlation between the observed and predicted phenotypes, improves from 0.15 (based on the genome alone) to 0.35 (based on the genome & exposome). We also show, using established theories, integrating genomic and exposomic data is essential to attaining a clinically meaningful level of prediction accuracy for disease traits. In conclusion, the genomic and exposomic effects can contribute to phenotypic variation via their latent relationships, i.e. genome-exposome correlation, and gxe and exe interactions, and modelling these effects has a great potential to improve phenotypic prediction accuracy and thus holds a great promise for future clinical practice.

## Introduction

Both genetic and environmental factors underlie phenotypic variance of complex traits. Understanding the influences of these factors not only helps explain why individuals differ from one another in phenotypes but also helps predict future phenotypes, such as disease diagnoses. The proliferation of genotypic data in the past decades, along with developments in relevant analytic tools, have already contributed a great deal to understanding phenotypic variations of complex traits^1-9^, and enabled phenotypic predictions at a level of accuracy for potential use in clinical settings^10-12^. However, these understandings and predictions are bounded by the heritability of the traits, and for many complex traits, large phenotypic variation remains unexplained, suggesting substantial environmental contributions to phenotypic variance.

Complementary to the genome, the concept of exposome has been proposed to capture the totality of human environmental exposures, encompassing external as well as internal environments over the lifetime of a given individual^13-15^. Similar to genotypes, exposomic variables are standardised across cohorts^16^. Since the inception of the concept, considerable efforts have been made to assess and characterise the exposome^17^. For example, the Human Early-Life Exposome project is a European collaborative effort established to characterize the early-life exposome which includes all environmental hazards that mothers and children are exposed to^18^. Despite the progress in the construction of the exposome, few analytic tools exist to date that can integrate genomic and exposomic data for complex trait analyses. We hypothesize that exposomic variables do not only affect phenotypes on their own but also interact among each other^19,20^ and with genotypes^20,21^. In addition, the estimation of exposomic effects and genomic effects on phenotypes could be biased, if these effects are correlated but the estimation model assumes otherwise^22^. Hence, tools that integrate genomic and exposomic data are required to capture variance as well as covariance components of phenotypes.

Here we propose a versatile linear mixed model that fulfils these requirements. The proposed approach jointly models the random effects of the genome and exposome and can be extended to capture genome-exposome and exposome-exposome interactions and genome-exposome correlations in the phenotypic analysis of a complex trait. It also allows us to model exposomic effects modulated by one or a few specific environmental variables. We demonstrate the proposed approach using traits from the UK biobank with 11 complex traits and 28 lifestyle exposures that were measured using a standard protocol.

## Results

### Method overview

We used a novel linear mixed model (LMM) to jointly model the effects of the genome and exposome on the phenotypes of a complex trait. The exposome here is restricted to 28 lifestyle exposures that were measured using a standard protocol (see Methods). Our model has three key features. First, it allows estimation of the correlation between genomic and exposomic effects, relaxing the assumption of independence between those effects as in a conventional LMM^22^. Second, the model can capture both additive and non-additive effects of the exposome and genome, i.e. pair-wise interactions between exposomic variables (exe interactions; e.g.^19^) and interactions between exposomic variables and genotypes (i.e., gxe interactions; e.g.^21^). Third, the model can handle correlated exposomic variables (see Methods & Supplementary Note 1) that may cause biased variance estimations of exposomic variables (e.g.^20^).

To illustrate the use of the model with real data, we selected 11 complex traits from the UK Biobank with heritability estimates above 0.05, including BMI, sitting height and years of education etc. (https://nealelab.github.io/UKBB_ldsc/), along with 28 lifestyle variables, including alcohol use, smoking, physical activity and dietary composition (see Methods for a detailed description). We performed the following analyses. First, for each trait, we used various models to estimate variance components of the additive and non-additive effects of the exposome and genome, including exe interactions and gxe interactions. The significance of the variance components was determined through a series of model comparisons using likelihood ratio tests (Table 1). Second, we extended the proposed model to examine the extent to which exposomic effects are modulated by covariates such as age, sex and socio-economic status (i.e., exc interactions). Third, we used 5-fold cross validation to show that the prediction accuracy increased significantly after accounting for the exposomic effects and exe interactions. Finally, we explored the potential clinical use of the proposed integrative analysis of genomic and exposomic data, by projecting its prediction accuracy for a disease trait in terms of the area under the receiver operating characteristic curve (AUC). The projection was based on well-established theories^23-30^ that express AUC as a function of sample size, proportions of variance explained by genomic and exposomic effects and the population prevalence of the disease.

**Table 1.**
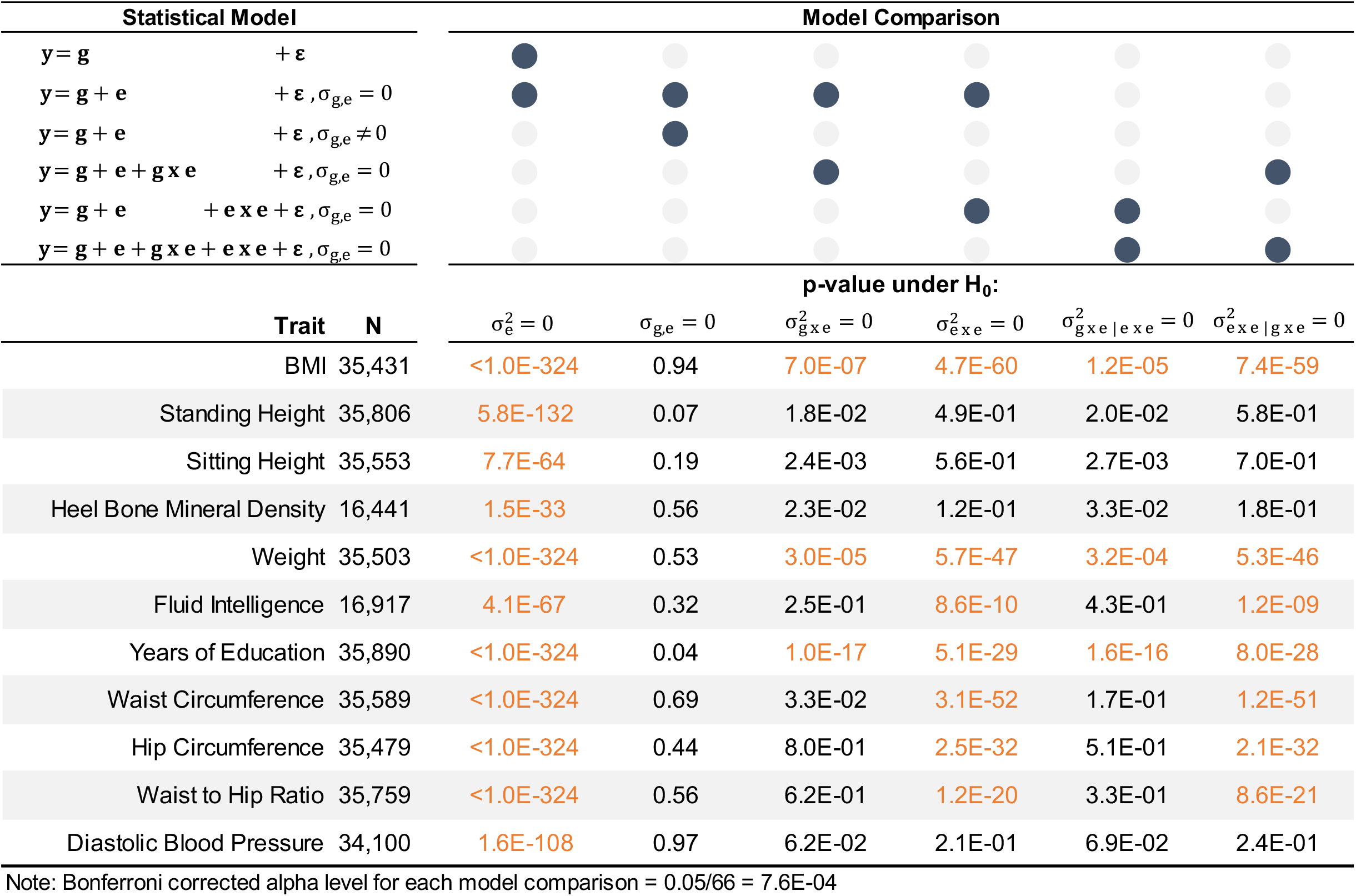
P-values for variance-covariance components of phenotypes of selected traits

### Exposomic effects on phenotypes

In line with previous estimation (https://nealelab.github.io/UKBB_ldsc/), we found significant SNP-based heritability for all selected traits, with estimates ranging between 0.08 (years of education) and 0.52 (standing height; Figure 1). We detected robust additive effects of the lifestyle-exposome on phenotypes of all traits (see Figure 1 for e and Table 1 for p-values under 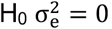). The magnitude of these additive effects, however, varied across traits. For example, the exposome accounted for 8.5% of the phenotypic variance of waist circumference, but less than 2.5 % for height, standing height, heel bone mineral density and fluid intelligence. Importantly, the additive exposomic effects were mostly uncorrelated with the genetic effects (see Table 1 for p-values under H_0_ σ_g,e_ = 0; see Supplementary Table 1 for covariance estimates), which was notably different from the genome-transcriptome correlation^22^.

**Figure 1.**
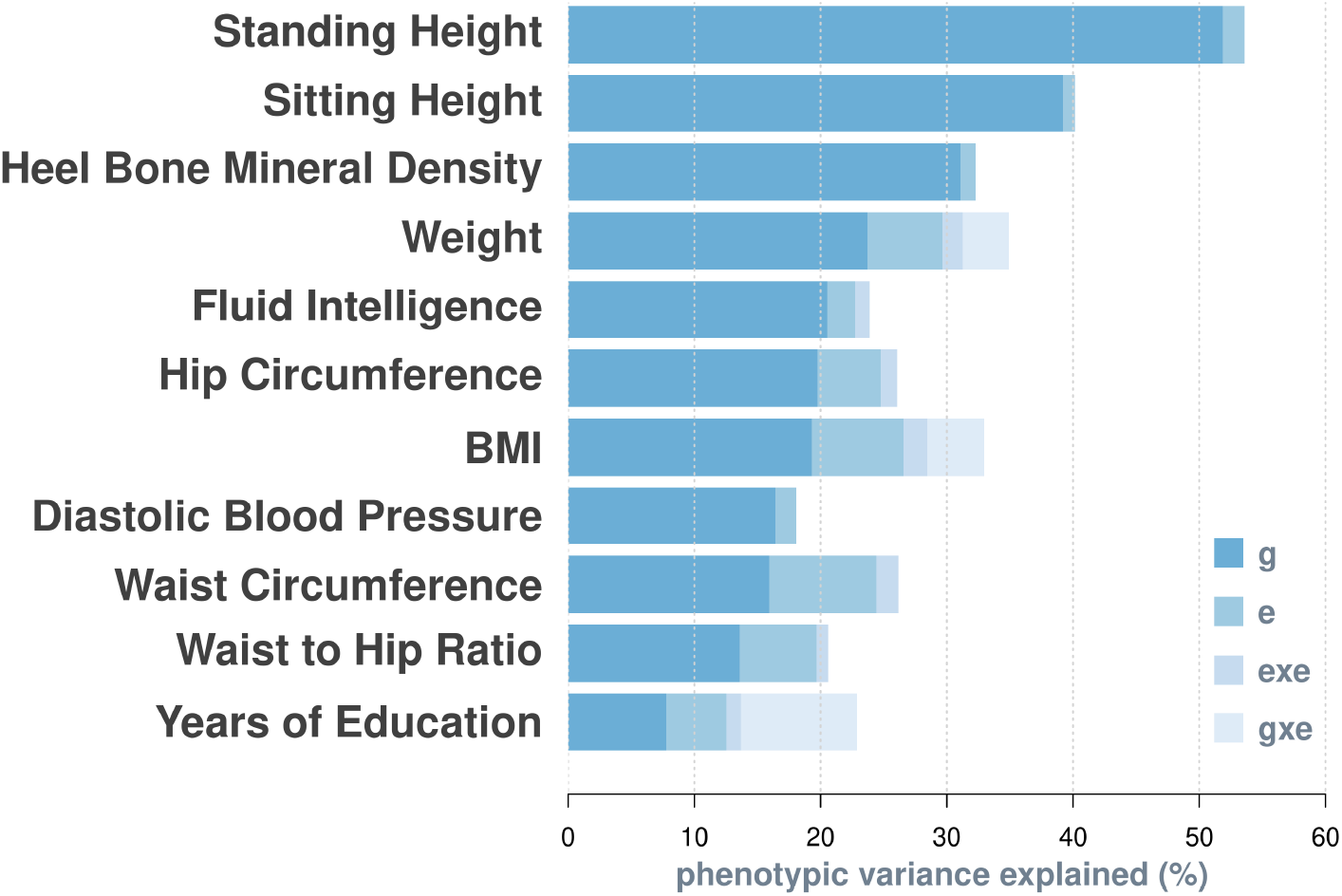
Breakdown of phenotypic variance by the model with the best fit. The best model for each trait is derived from model comparisons shown in Table 1. g = additive genetic effects on phenotypes; e = additive effects of exposomic variables; exe = interaction effects between exposomic variables; gxe = interaction effects between genotypes and exposomic variables. Variance components are expressed as percentage of total phenotypic variance.

The estimated variance component of non-additive effects of the lifestyle-exposome (exe) was highly significant for 7 out 11 traits (Table 1), although they only account for ∼ 1% to 2% of phenotypic variance (See Figure 1 & Supplementary Table 2). By contrast, significant gxe interactions are only evident for BMI, weight and years of education (Table 1), but they could account for up to 9% of total phenotypic variance (years of education; Figure 1 & Supplementary Table 2). The low presence of gxe signals is probably due to relatively low power of detecting gxe interactions, which is caused by a large number of pairs of gxe interaction terms to be estimated in the model, i.e. 28 (number of exposomic variables) x 1.3 million (number of SNPs) in this study. In addition, the identified gxe and exe interactions are orthogonal to each other. This is evidenced by that both gxe and exe remained significant when being jointly modelled (see p-values under H_0_ σ_gxe|exe_ = 0 and under H_0_ σ_exe|gxe_ = 0).

**Table 2.** Model equations and their assumed sample variance-covariance matrices.

| Model Equation | Matrix Notion | Sample Variance-Covariance Matrix |
| --- | --- | --- |
| For individual $i = 1, 2, \dots, n$,<br>**1** $y_i = \mu + \underbrace{\sum_{j=1}^{m_1} a_{ij} \alpha_j}_{g_i} + \varepsilon_i$<br>where $y_i$ is the phenotype, $\mu$ is the grand mean, $a_{ij}$ is the SNP genotype at locus $j$, $m_1$ is the total number of SNPs, $\alpha_j$ is the random effect of the SNP that is assumed to be normal with mean zero and variance $\sigma_g^2/m_1$, and $\varepsilon_i$ is the residual assumed to be normal with mean zero and variance $\sigma_\varepsilon^2$. | For $\mathbf{y} = (y_1, y_2, \dots, y_n)$,<br>$\mathbf{y} = \mu \mathbf{1}_n + \mathbf{g} + \boldsymbol{\varepsilon}$<br>where $\mathbf{g} = \mathbf{A}\boldsymbol{\alpha}^t = \begin{pmatrix} a_{11} & \cdots & a_{1m_1} \\ \vdots & \ddots & \vdots \\ a_{n1} & \cdots & a_{nm_1} \end{pmatrix} \begin{pmatrix} \alpha_1 \\ \vdots \\ \alpha_{m_1} \end{pmatrix}$ | $\sigma_g^2 \underbrace{\mathbf{A}\mathbf{A}^t/m_1}_{\mathbf{G}} + \sigma_\varepsilon^2 \mathbf{I}$<br>where $\mathbf{I}$ is the $n \times n$ identity matrix. |
| **2a** $y_i = \mu + g_i + \underbrace{\sum_{k=1}^{m_2} b_{ik} \beta_k}_{e_i} + \varepsilon_i$<br>where $b_k$ is the $k$th exposomic variable, $m_2$ is the total number of exposomic variables, and $\beta_k$ is the random effect of the exposomic variable that is assumed to be normal with mean zero and variance $\sigma_e^2/m_2$. To avoid estimation bias due to multicollinearity, $b_k$ is transformed using a principal component analysis (see Methods). | $\mathbf{y} = \mu \mathbf{1}_n + \mathbf{g} + \mathbf{e} + \boldsymbol{\varepsilon}$<br>where $\mathbf{e} = \mathbf{B}\boldsymbol{\beta}^t = \begin{pmatrix} b_{11} & \cdots & b_{1m_2} \\ \vdots & \ddots & \vdots \\ b_{n1} & \cdots & b_{nm_2} \end{pmatrix} \begin{pmatrix} \beta_1 \\ \vdots \\ \beta_{m_2} \end{pmatrix}$ | $\sigma_g^2 \mathbf{G} + \sigma_e^2 \underbrace{\mathbf{B}\mathbf{B}^t/m_2}_{\mathbf{E}} + \sigma_\varepsilon^2 \mathbf{I}$ |
| **2b** $y_i = \mu + g_i + e_i + \varepsilon_i$ | $\mathbf{y} = \mu \mathbf{1}_n + \mathbf{g} + \mathbf{e} + \boldsymbol{\varepsilon}$ | $\sigma_g^2 \mathbf{G} + \sigma_e^2 \mathbf{E} + \left[ \sqrt{\mathbf{G}} + \sqrt{\mathbf{E}^t} + \left( \sqrt{\mathbf{G}} + \sqrt{\mathbf{E}^t} \right)^t \right] \sigma_{ge} + \sigma_\varepsilon^2 \mathbf{I}$<br>where $\sqrt{\mathbf{G}}$ and $\sqrt{\mathbf{E}^t}$ are the Cholesky decompositions of $\mathbf{G}$ and $\mathbf{E}^t$, respectively, and $\sigma_{ge}$ is the covariance between $\mathbf{g}$ and $\mathbf{e}$. |
| **3** $y_i = \mu + g_i + e_i + \underbrace{\sum_{q=1}^Q c_{iq} \gamma_q}_{\mathbf{g} \times \mathbf{e}_i} + \varepsilon_i$<br>where $c_q$ is the $q$th pairwise interaction term between SNP genotypes and exposomic variables, and $\gamma_q$ is the effect of the $q$th interaction term. $\gamma_q$ is assumed to be normally distributed with mean zero and variance $\sigma_{g \times e}^2/Q$, and $Q$ is the total number of interaction terms ($Q = m_1 m_2$). | $\mathbf{y} = \mu \mathbf{1}_n + \mathbf{g} + \mathbf{e} + \mathbf{g} \times \mathbf{e} + \boldsymbol{\varepsilon}$<br>where $\mathbf{g} \times \mathbf{e} = \mathbf{C}\boldsymbol{\gamma}^t = \begin{pmatrix} c_{11} & \cdots & c_{1Q} \\ \vdots & \ddots & \vdots \\ c_{n1} & \cdots & c_{nQ} \end{pmatrix} \begin{pmatrix} \gamma_1 \\ \vdots \\ \gamma_Q \end{pmatrix}$, and $\mathbf{C}$ can be derived using the following pseudo-code with $\mathbf{A} = [\mathbf{a}_1 \ \cdots \ \mathbf{a}_{m_1}]$; $\mathbf{B} = [\mathbf{b}_1 \ \cdots \ \mathbf{b}_{m_2}]$; $\mathbf{C} = [\mathbf{c}_1 \ \cdots \ \mathbf{c}_Q]$, and $q = 1, 2 \dots Q$.<br>for $i = 1$ to $m_1$ {<br>for $j = 1$ to $m_2$ {<br>$\mathbf{c}_q = \mathbf{a}_i \otimes \mathbf{b}_j$ } } } | $\sigma_g^2 \mathbf{G} + \sigma_e^2 \mathbf{E} + \sigma_{g \times e}^2 \boldsymbol{\Gamma} + \sigma_\varepsilon^2 \mathbf{I}$<br>where $\boldsymbol{\Gamma}$ is a $n \times n$ matrix derived by the Hadamard product of $\mathbf{G}$ and $\mathbf{E}$, i.e., $\mathbf{G} \otimes \mathbf{E}$. |
| **4** $y_i = \mu + g_i + e_i + \underbrace{\sum_{p=1}^P x_{ip} \theta_p}_{\mathbf{e} \times \mathbf{e}_i} + \varepsilon_i$<br>where $x_p$ is the $p$th pairwise interaction term between exposomic variables, and when the two exposomic variables are identical, the interaction term becomes the quadratic term of the exposomic variable; $\theta_p$ is the effect of the $p$th interaction term and is assumed to be normally distributed with mean zero and variance $\sigma_{\mathbf{e} \times \mathbf{e}}^2/P$, and $P$ is the total number of interaction terms ($P = m_2(m_2 + 1)/2$). To avoid estimation bias due to multicollinearity, $x_p$ is transformed using a principal component analysis (see Methods). | $\mathbf{y} = \mu \mathbf{1}_n + \mathbf{g} + \mathbf{e} + \mathbf{e} \times \mathbf{e} + \boldsymbol{\varepsilon}$<br>where $\mathbf{e} \times \mathbf{e} = \mathbf{X}\boldsymbol{\theta}^t = \begin{pmatrix} x_{11} & \cdots & x_{1P} \\ \vdots & \ddots & \vdots \\ x_{n1} & \cdots & x_{nP} \end{pmatrix} \begin{pmatrix} \theta_1 \\ \vdots \\ \theta_P \end{pmatrix}$,<br>and $\mathbf{X}$ can be derived using the following pseudo-code with $\mathbf{B} = [\mathbf{b}_1 \ \cdots \ \mathbf{b}_{m_2}]$; $\mathbf{X} = [\mathbf{x}_1 \ \cdots \ \mathbf{x}_P]$, and $p = 1, 2 \dots P$.<br>for $i = 1$ to $m_2${<br>for $j = i$ to $m_2${<br>$\mathbf{x}_p = \mathbf{b}_i \otimes \mathbf{b}_j$ } } } | $\sigma_g^2 \mathbf{G} + \sigma_e^2 \mathbf{E} + \sigma_{\mathbf{e} \times \mathbf{e}}^2 \underbrace{\mathbf{X}\mathbf{X}^t/P}_{\boldsymbol{\Theta}} + \sigma_\varepsilon^2 \mathbf{I}$ |
| **5** $y_i = \mu + g_i + e_i + g \times e_i + e \times e_i + \varepsilon_i$ | $\mathbf{y} = \mu \mathbf{1}_n + \mathbf{g} + \mathbf{e} + \mathbf{g} \times \mathbf{e} + \mathbf{e} \times \mathbf{e} + \boldsymbol{\varepsilon}$ | $\sigma_g^2 \mathbf{G} + \sigma_e^2 \mathbf{E} + \sigma_{\mathbf{g} \times \mathbf{e}}^2 \boldsymbol{\Gamma} + \sigma_{\mathbf{e} \times \mathbf{e}}^2 \boldsymbol{\Theta} + \sigma_\varepsilon^2 \mathbf{I}$ |
| **6** $y_i = \mu + g_i + e_i + \sum_{l=1}^L c_{il} \underbrace{\sum_{k=1}^{m_2} b_{ik} \lambda_{kl}}_{\mathbf{e}_{li}} + \varepsilon_i$<br>where $\lambda_{kl}$ is the random effect of $k$th exposomic variable, $b_k$, modulated by the $l$th covariate $c_l$. $\lambda_{kl}$ is assumed to be normally distributed with mean zero and variance $\sigma_{e_l}^2/m_2$ | $\mathbf{y} = \mu \mathbf{1}_n + \mathbf{g} + \mathbf{e} + \sum_{l=1}^L \mathbf{e} \times \mathbf{c}_l + \boldsymbol{\varepsilon}$<br>where $\mathbf{e} \times \mathbf{c}_l$ is a $n \times 1$ vector that can be derived by $\mathbf{e}_l \otimes \mathbf{c}_l$, and $\mathbf{e}_l = \begin{pmatrix} b_{11} & \cdots & b_{1m_2} \\ \vdots & \ddots & \vdots \\ b_{n1} & \cdots & b_{nm_2} \end{pmatrix} \begin{pmatrix} \lambda_{1l} \\ \vdots \\ \lambda_{m_2 l} \end{pmatrix}$ | $\sigma_g^2 \mathbf{G} + \mathbf{E} \otimes (\boldsymbol{\Phi} \mathbf{K} \boldsymbol{\Phi}^t) + \sigma_\varepsilon^2 \mathbf{I}$<br>where $\boldsymbol{\Phi} = (\mathbf{1}_n \ \mathbf{c}_1 \ \mathbf{c}_2 \ \cdots \ \mathbf{c}_L)$ and $\mathbf{K} = \begin{pmatrix} \sigma_{e_0}^2 & \cdots & \sigma_{e_0 e_L} \\ \vdots & \ddots & \vdots \\ \sigma_{e_0 e_L} & \cdots & \sigma_{e_L}^2 \end{pmatrix}$. |

By extending the proposed model to a reaction normal model (RNM; see Methods), we examined whether the additive exposomic effects on phenotype vary depending on specific covariates, which would be evidenced by the presence of significant exc interactions. Using single-covariate RNMs, we identified several significant exc interactions (Supplementary Table 3), noting that most covariates are lifestyle related, which are in line with the exe interactions found above. For each trait, we then fitted an RNM model that simultaneously includes all significant exc interactions identified from single-covariate RNM analyses. The variance estimates of exc interactions from the joint analyses are presented in Supplementary Table 7.

**Table 3.**
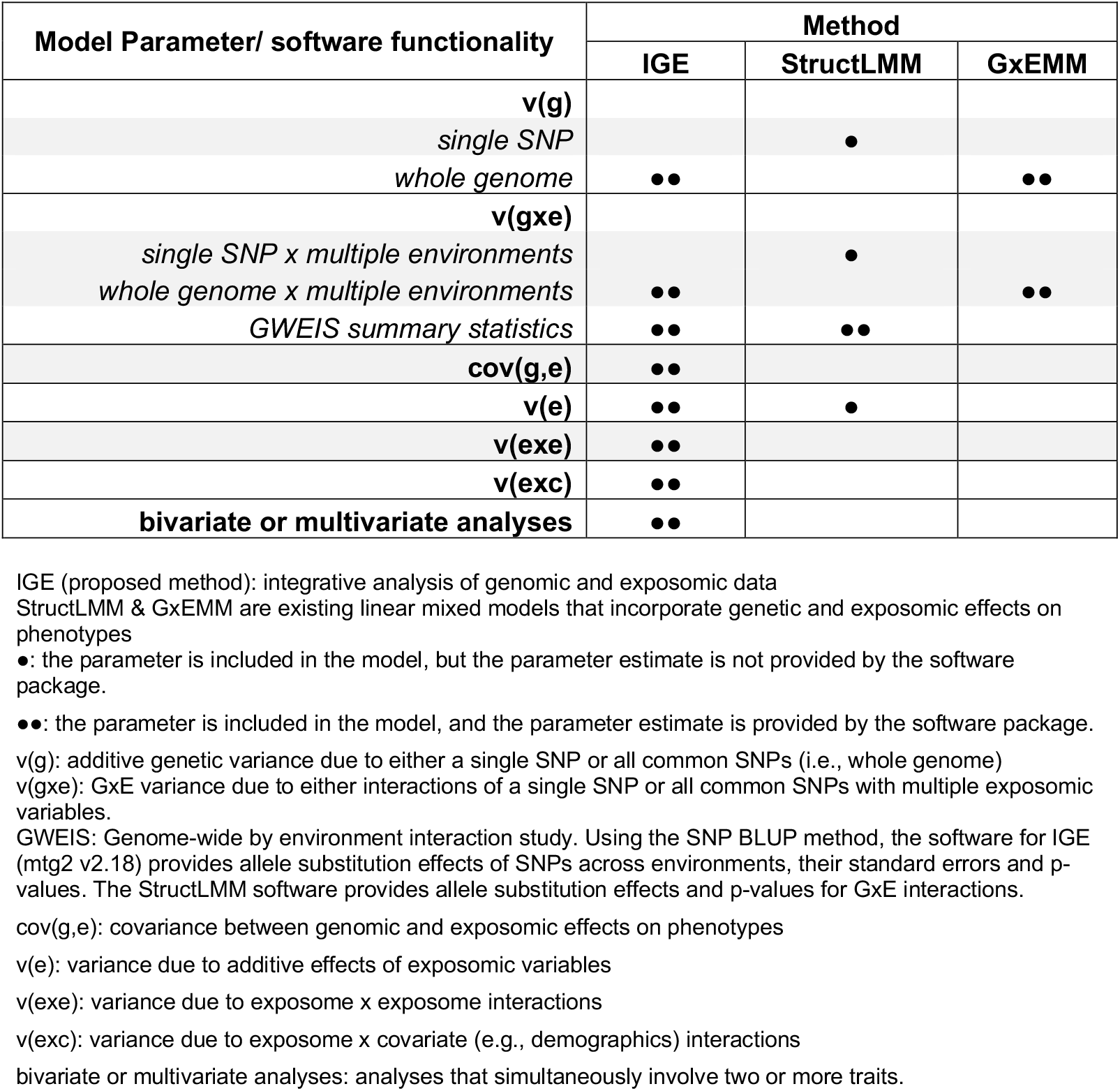
Comparisons of methods (software packages) on the genomic and exposomic analysis of complex traits

It is important to note that the estimation of exposomic effects is sensitive to the correlation structure of exposomic variables. Specifically, multicollinearity between exposomic variables would bias the estimate of 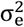 (Supplemental Note 1); and by extension, correlated exe interaction terms and gxe interaction terms (Equations 3 & 4 in Table 2) would bias the estimates of 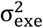 and 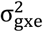, unless their true values are small (e.g., 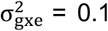 and 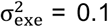in our simulations). Without knowing the true values of variance components, transforming exposomic variables and interaction terms using a principal component analysis (see Methods & Supplemental Note 1) seems necessary prior to model fitting in order to avoid estimation bias due to multicollinearity. While transforming the exposomic variables and the exe interaction terms are computationally trivial, transforming the gxe interaction terms is computationally infeasible (28 x 1.3 million variables). Nonetheless, the variance of gxe interactions is small in general, suggesting that using the gxe interaction terms without the transformation (i.e., derived from $⨂ & in Equation 3 of Table 2) is generally free from the estimation bias due to multicollinearity. Note that the largest variance estimate of gxe interactions in this study is ∼0.09.

### Validation of exposomic effects

Using 5-fold cross-validation, we found that both additive (e) and non-additive effects (exe) of the exposome, which were significant in the discovery dataset, could improve the phenotypic prediction accuracy in the target dataset. In general, the larger the variance estimates, the greater the prediction improvements (Figures 2a & 2b), which indicates that the additive effects of the exposomic variables and exe interactions are genuine. Similarly, we also validated the exposomic effects modulated by specific covariates, by showing that the larger the total variance estimates of exc interactions, the greater the improvement of predication accuracy (Figure 3). The validated exc interactions would in part explain the phenotypic variance due to residual x covariate interactions found in our previous studies^31-33^.

**Figure 2.**
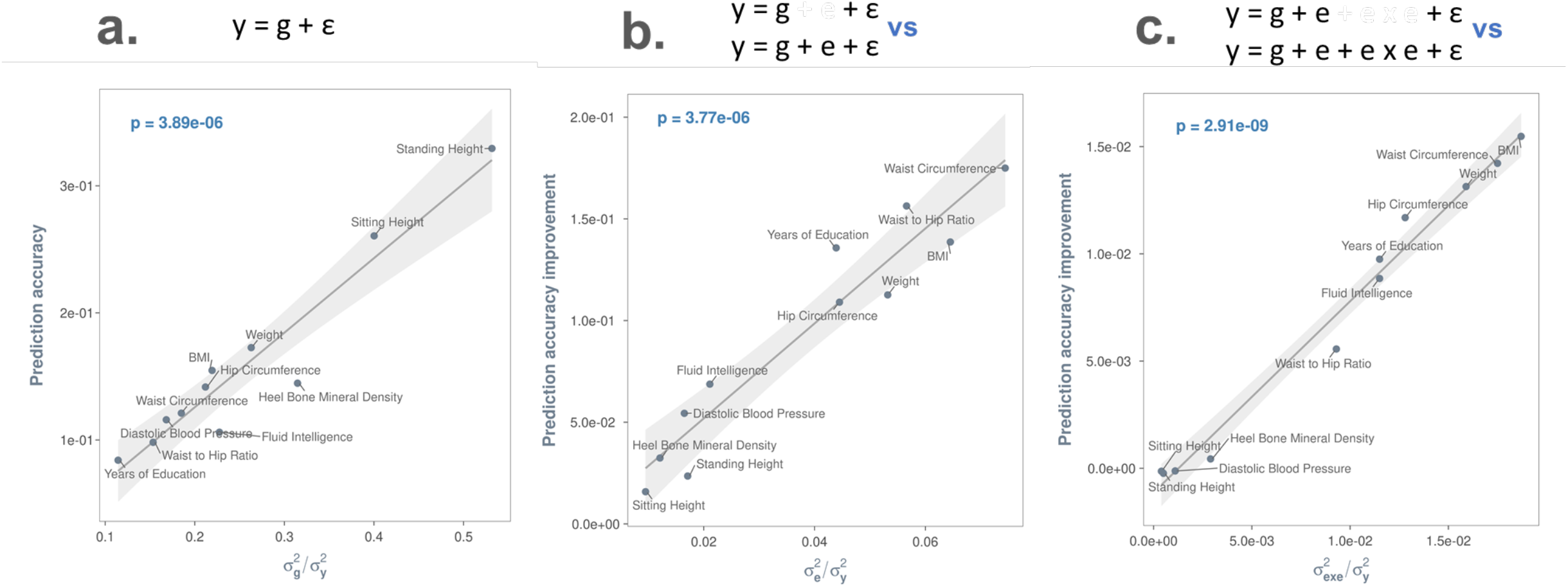
Exposomic variables contribute to phenotypic variance and improve phenotypic prediction accuracy. The prediction accuracy of a given model was computed using the Pearson’s correlation coefficient between the observed and the predicted by the model. For all panels, the least squares line with 95% confidence band is based on a linear model that regressed either prediction accuracies (panel a) or predication accuracy improvements (panels b-c) by a model on variance component estimates of the model. The p-value is for the t-test statistic (df=7) under the null hypothesis that the slope of the regression line is zero. 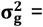 phenotypic variance explained by additive effects of the genome; 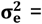 phenotypic variance explained by additive effects of the exposome; 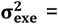 phenotypic variance explained by exposome-by-exposome interactions; and 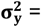 = total phenotypic variance. Panel a. Phenotypic prediction accuracies by the baseline model that uses genomic data alone, i.e., y = g+ε, where g = phenotypic effects of the genome and ε = residuals. The larger the genetic variance, the greater the prediction accuracy. Panel b. Additive effects of the exposomic variables (i.e., e) contribute to phenotypic variance and improve phenotypic prediction accuracy. The greater the additive effects, the larger the gain in phenotypic prediction accuracy. A prediction accuracy improvement (on the y-axis) was derived by subtracting the prediction accuracy of the model y = g+ε from that of the model y = g+e+ε. Panel c. Exposome-by-exposome interactions (i.e., exe interactions) contribute to phenotypic variance and further improve phenotypic prediction accuracy. The greater the variance estimate of exe interactions, the larger the gain in phenotypic prediction accuracy. A prediction accuracy improvement (on the y-axis) was derived by subtracting the prediction accuracy of the model y = g+e+ε from that of the model y = g+e+exe+ε.

**Figure 3.**
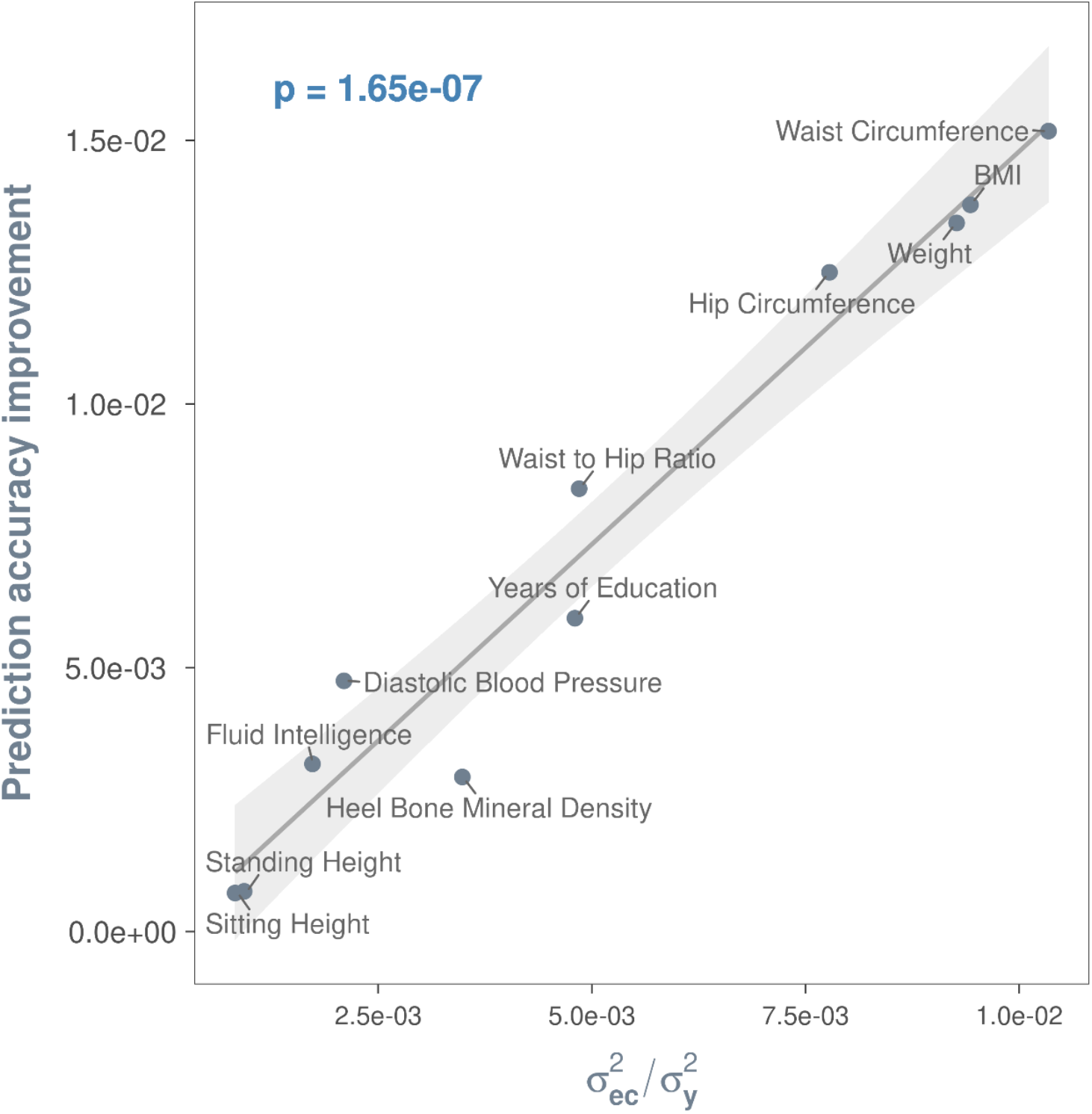
Positive relationship between phenotypic variance explained by exposome-by-covariate (exc) interaction effects and prediction accuracy improvement. Prediction accuracy improvement is computed by subtracting the prediction accuracy of the model y = g + e + ε from that of a model with multiple covariates (see Equation 6 of Table 2) that are shown to interact with the exposome in univariate exc interaction analyses. The least squares line with 95% confidence band is based on a linear model that regressed prediction accuracy improvement on phenotypic variance explained by exc interactions. The p-value is for the t-test statistic (df=7) under the null hypothesis that the slope of the regression line is zero. Significant covariates included for each trait can be found in Supplementary Table 3.

By contrast, although gxe interactions contribute to the phenotypic variance of BMI, weight and years of eduation (Table 1), the contribution did not lead to significant gains in phenotypic prediction accuracy (Supplementary Figure 1). This was most likely due to a lack of power. i.e. the size of discovery samples was insufficient to accurately estimate an extremely large number of parameters, i.e., best linear unbiased prediction (BLUPs) of gxe interaction effects^23,27,28,34^. This is further verified using simulations (see Supplementary Note 2 and Supplementary Figure 2).

Given the sample sizes of the discovery data sets (∼28,000), the prediction accuracies of the model y = g + ε for the selected traits are only between 1/3 and 1/2 of the theoretical maximums (i.e., square root of heritability; Supplementary Figure 3). They can improve, in theory, by increasing the sample size of discovery sets (Supplementary Figure 3); or, as shown in the above, by accounting for the additive effects of the exposome and exe interactions (Figures 2b & 2c). To examine prediction efficiency of the latter, we projected the observed prediction accuracies of the models y = g + e + ε and y = g + e + exe + ε onto the theoretical trajectory of prediction accuracies of the model y = g + ε as a function of the sample sizes of discovery datasets (Supplementary Figure 3). As such, the use of exposomic information could improve phenotypic prediction accuracy to the same extent as a 1.2 to 14-fold increase in sample size, depending on the significance of the exposomic effects and their interactions (Figure 4). Given the substantial costs and efforts associated with increasing sample size, the improved predictive accuracy by the models y = g + e + ε and y = g + e + exe + ε are considerable, despite the fact that the proportion of phenotypic variance explained by the exposome is small (see the x-axis of Figures 2b & 2c).

**Figure 4.**
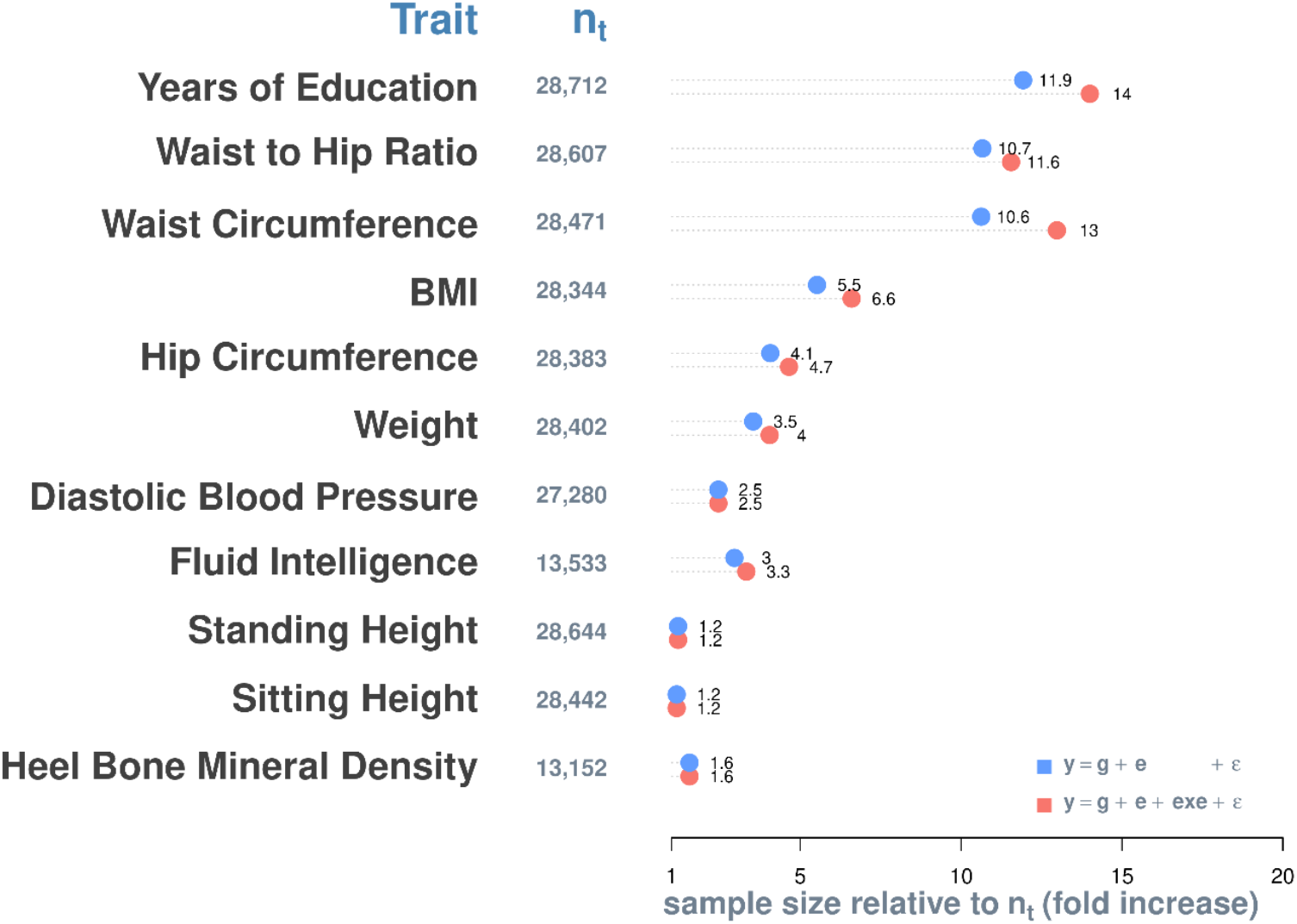
Additional sample size required for the model y = g + ε to achieve the same level of prediction accuracy as y = g + e + ε (blue) and y = g + e + exe + ε (red). n_t_ = sample size of training (or discovery) datasets.

### Quantification of clinical relevance

We quantified the clinical relevance of the proposed model by exploring its prediction accuracy for quantitative traits and disease traits. For quantitative traits, we expressed the prediction accuracy of the model y = g + e + ε (i.e., correlation coefficient between the true and predicted phenotypes) as a function of the sample size of the discovery dataset, variances explained by the genome and exposome, and effective numbers of (independent) SNPs and exposomic variables (see Methods), using previous theoretical derivations^27-30,34^. Based on the derived expression (Equation 6), we computed the expected prediction accuracies for the quantitative traits used in this study and found that they agreed well with the observed prediction accuracies from the 5-fold cross validation (Supplementary Figure 4). We then extended the derived expression to disease traits in terms of the area under the operative characteristic curve (AUC; see Equation 10 in Methods for details) using well-established theories^23-26^. AUC is a gold-standard measure used to evaluate how well a prediction model discriminates diseased from non-diseased individuals. An AUC between 0.7 to 0.8 is considered acceptable, 0.8 to 0.9 excellent, and above 0.9 outstanding^35^. Figure 5 shows the expected AUC of the proposed integrative analysis of genomic and exposomic data for disease traits of different values of population prevalence (k), assuming different amounts of variance (on the liability scale) explained by the genome and exposome and discovery sample sizes. For simplicity, we use 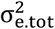 to denote the total variance in disease liability explained by additive effects of the exposome and exe interactions as a whole.

**Figure 5.**
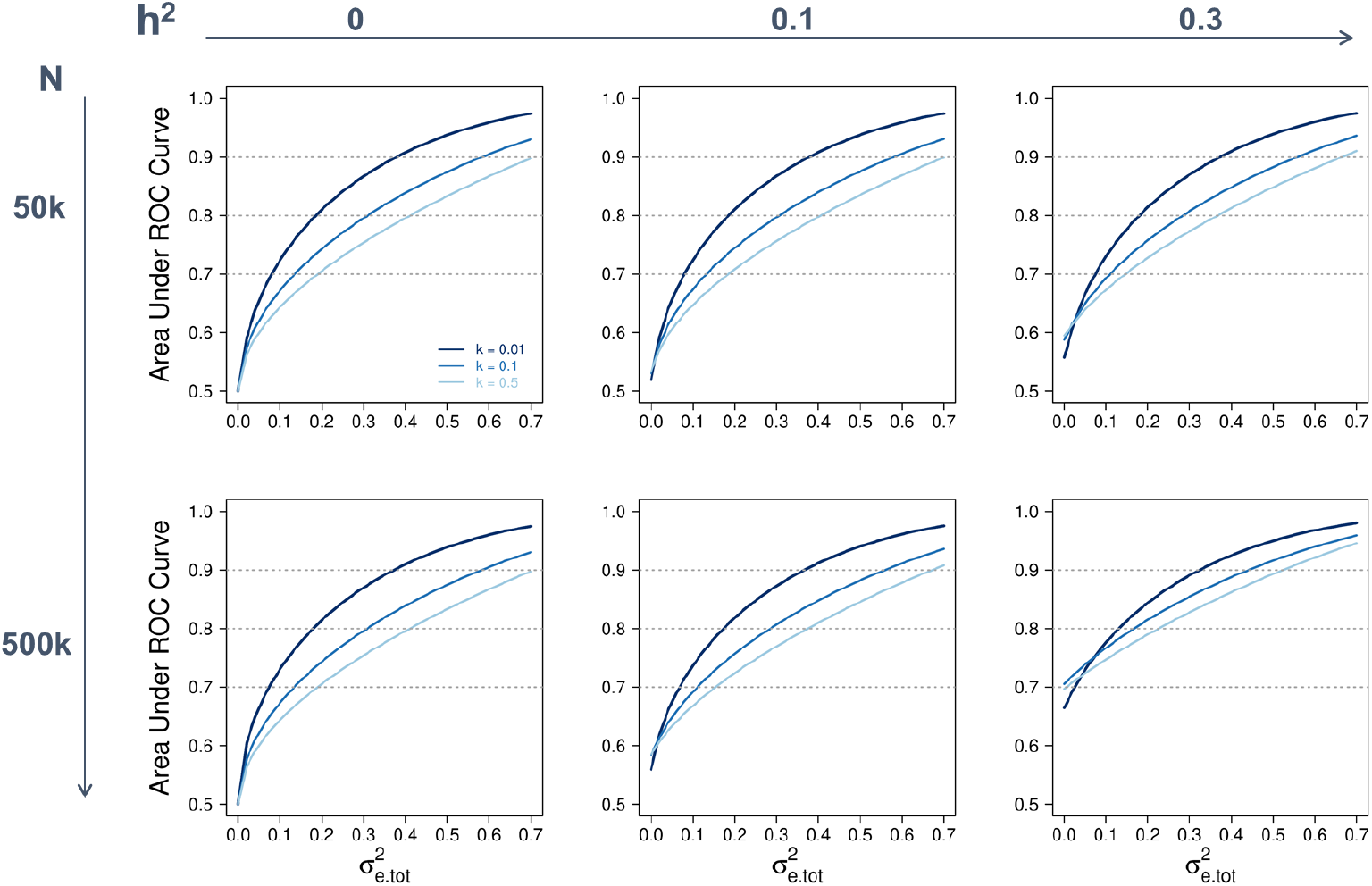
Expected prediction accuracy of the proposed integrative analysis of genetic and exposomic data for disease traits of different prevalence (k) and heritability (h^2^) at varying levels of total variance explained by the exposome 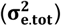 and sample size of the discovery dataset (N). Diseases are assumed to have a liability of mean zero and variance 1, and both h^2^ and 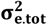 are on the disease liability scale. Prediction accuracy is measured using the area under the receiver operating characteristic (ROC) curve, with 0.7 to 0.8 generally being considered acceptable, 0.8 to 0.9 excellent, and above 0.9 outstanding. The assumed effective number of chromosome segments and the number of exposomic variables are 50,000 and 28, respectively, which are based on the genomic and exposomic data used in this study. However, varying the number of exposomic variables from 28 to 100 does not have a notable effect on the expected area under the ROC curve.

When setting 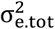 to 0—that is, using no exposomic information at all—varying the heritability of disease liability h^2^ from 0 to 0.3 improves AUC from 0.5 to ∼ 0.6 when the sample size of the discovery set is 50k. This is in contrast to a 2-fold improvement, from 0.5 to ∼ 0.7, when the sample size is 500k. Thus, genomic prediction accuracy heavily relies on sample size, such that for a disease trait with a moderate heritability, a clinically meaningful level of accuracy (AUC >=0.7) may not be attainable unless the sample size of the discovery dataset is substantially large (> = 500k). On the other hand, the benefit of using exposomic information to disease prediction can be realised with a relatively small discovery sample. This is evidenced by that when setting h^2^ to 0 (i.e., using no genomic information at all), increasing the value of 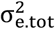 has the same effects on AUC whether using a discovery sample of 50k or 500k individuals. Notably, for a moderately heritable disease that affects 1% of the population, with a discovery dataset of 50k individuals, a collection of exposomic variables that together explains ∼30% of the variance in disease liability is sufficient to yield an AUC greater than 0.85 for the target sample (see area under ROC when h^2^ = 0.3, k = 0.01, 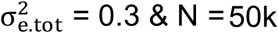 & N = 50k in Figure 5). Importantly, in all scenarios, AUC consistently improves with increasing 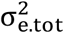. Thus, incorporating exposomic data is not only an efficient but also an effective way of improving prediction accuracy based on genomic data alone. Taken together, genomic prediction accuracy for disease traits is constrained by sample size; with a relatively small sample at hand, a desired level of prediction accuracy may only be achieved by combining genomic and exposomic information.

### Comparison with existing models

The key model parameters of the proposed integrative analysis of genomic and exposomic data (IGE) compared to existing linear mixed models that incorporate genetic and environmental effects on phenotypes are outlined in Table 3. In general, IGE offers thus far the most detailed partition of phenotypic variance.

Both IGE and GxEMM^36^ are whole-genome approaches to the estimation of heritability and g x e interactions, although IGE is considered more comprehensive and versatile, which models variances explained by additive effects of exposomic variables, by exposome x exposome interactions, and by exposome x covariate (such as demographics) interactions; and covariance between genetic effects and exposomic effects (Table 3). Further, bivariate or multivariate IGE (i.e., simultaneously including two or more traits) can be feasibly performed using mtg2 version 2.18 (https://sites.google.com/site/honglee0707/mtg2).

In contrast, StructLMM has been developed primarily for a genome-wide by environment interaction study (GWEIS)^20^ that examines one SNP at a time with a focus on association tests (providing p-values) for GxE interactions between the SNP genotypes and multiple exposomic variables. Using the well-established SNP BLUP method^2,37,38^, IGE can also provide GWEIS summary statistics, including estimated allele substitution effects of all SNPs across environments, their standard errors and p-values. Note that SNP BLUP implemented in IGE can model all SNP jointly (a whole-genome approach). Nonetheless, one of the main scopes of this study is to provide unbiased estimates of exposomic variances, e.g. v(e) 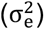 that is common to both StructLMM and IGE (Table 3 & Supplementary Note 3). Importantly, correlated exposomic variables would cause biased estimation of 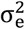 (Supplementary Table 4) unless they are transformed to independent variables via a principal component analysis (Methods). To our knowledge, this transformation has not yet been implemented in any existing methods including StructLMM. Using results from simulations, we show that 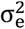 estimates by StructLMM are prone to bias due to correlated environments (Supplementary Table 4). The other model parameters such as v(exe), v(exc) and cov(g, e) cannot be estimated by StructLMM (Table 3).

## Discussion

Using our approach, we demonstrate the importance of the exposome for understanding individual differences in phenotypes. Although the ‘exposome’ constructed in this study comprises only 28 lifestyle factors, when integrated with genomic data, it explained between 2% to 10% additional phenotypic variance and significantly improved phenotypic prediction accuracy to a level equivalent to a 1.2 to 14-fold increase in sample size. The additional phenotypic variance is not only from additive effects of the exposome but also from its non-additive effects (exe) and genome-exposome interactions (gxe). We expect that as the construction of the exposome continues to progress, more phenotypic variance will be explained and greater improvements in phenotypic prediction accuracy will be gained. This would be particularly promising for phenotypic analysis and prediction of traits with small to little heritability component, such as ovarian and colorectal cancer^39^.

We noted that when exposomic variables are correlated, the variance estimate of additive effects of exposomic variables is biased unless these variables are transformed using a principal component analysis (i.e. **E** in Table 2 should be based on transformed variables). By extension, this would apply to exe interaction terms and gxe interactions terms, unless the proportions of phenotypic variance explained by these interaction effects are small (<10%), as shown in our simulations. These observations have important implications for modelling environmental effects in LMMs. Recently, Moore et al.^20^ proposed the structured linear mixed model (StructLMM) that incorporates random effects of multiple environments in order to study the interactions between these environments and genotypes of a single SNP (i.e., gxe interactions). However, the environmental variables in StructLMM are not transformed, even though they are very likely correlated, which would have biased the variance estimate of environmental effects. Consequently, it remains uncertain the extent to which the estimation bias affects the StructLMM-based test statistics for detecting gxe interactions.

Depending on the research question at hand, the construction of the exposome may be guided by causal analyses. A meaningful exposome may only contain causal information. Examples may include lifestyles that potentially alter the molecular pathways or the pathogenesis of the main trait, or biomarkers that potentially explain possible molecular pathways underlying the phenotypes. As a contrast, in our BMI analysis, for example, it is not useful to include weight and height as part of the exposome, even though they would explain a large amount of phenotypic variance. This is because variations in these traits inform nothing other than the fact that they are correlated with the trait.

Heritability estimates were slightly reduced after including more variance components (result not shown). We considered two possibilities. First, the exposome may mediate part of additive genetic effects on phenotypes. For example, some SNPs affect smoking status, which in turn affect BMI. A model that simultaneously includes genetic and exposomic data would account for smoking status and their genetic effects, and hence gives arise to reduced heritability estimates. Second, there is a genuine correlation between exposomic and genomic effects in some latent mechanism. It is noted that there are marginally significant correlation estimates, which were not significant after Bonferroni correction. Such correlation may be because people who have similar genotypes may somehow share similar exposures i.e. genotype-environment correlation^40^.

In conclusion, the genomic and exposomic effects can contribute to phenotypic variation via their latent relationships, i.e. genome-exposome correlation, and gxe and exe interactions, for which our proposed method can provide reliable estimates. We show that the integrative analysis of genomic and exposomic data has a great potential for understanding genetic and environmental contributions to complex traits and for improving phenotypic prediction accuracy, and thus holds a great promise for future clinical practice.

## Methods

### Ethics statement

We used data from the UK Biobank (http://www.ukbiobank.ac.uk/) for our analyses. The UK Biobank’s scientific protocol has been reviewed and approved by the North West Multi-centre Research Ethics Committee (MREC), National Information Governance Board for Health & Social Care (NIGB), and Community Health Index Advisory Group (CHIAG). UK Biobank has obtained informed consent from all participants. Our access to the UK Biobank data was under the reference number 14575. The research ethics approval of the current study was obtained from the University of South Australia Human Research Ethics Committee.

### Genotype data

The UK Biobank contains health-related data from ∼ 500,000 participants aged between 40 and 69, who were recruited throughout the UK between 2006 and 2010^41^. Prior to data analysis, we applied stringent quality control to exclude unreliable genotypic data. We filtered SNPs with an INFO score (used to indicate the quality of genotype imputation) < 0.6, a MAF < 0.01, a Hardy-Weinberg equilibrium p-value <1e-4, or a call rate < 0.95. We then selected HapMap phase III SNPs, which are known to yield reliable and robust estimates of SNP-based heritability^42-44^, for downstream analyses. We filtered individuals who had a genotype-missing rate > 0.05, were non-white British ancestry, or had the first or second ancestry principal components outside six standard deviations of the population mean. We also applied quality control on the degree of relatedness between individuals by excluding one of any pair of individuals with a genomic relationship > 0.025. From the remaining individuals, we selected those who were included in both the first and second release of UK Biobank genotype data. Eventually, 408,218 individuals and 1,133,273 SNPs passed the quality control of genotype data.

### Phenotype data

We chose eleven UK Biobank traits available to us that have a heritability estimate (by an independent open source; https://nealelab.github.io/UKBB_ldsc/) greater than 0.05. These traits are standing height, sitting height, body mass index, heel bone mineral density, fluid intelligence, weight, waist circumference, hip circumference, waist-to-hip ratio, diastolic blood pressure and years of education.

Prior to model fitting, phenotypic data were prepared using R (v3.4.3) in three sequential steps: 1) adjustment for age, sex, birth year, social economic status (by Townsend Deprivation Index), population structure (by the first ten principal components of the genomic relationship matrix estimated using PLINK v1.9), assessment centre, and genotype batch using linear regression; 2) standardization; and 3) removal of data points outside +/-3 standard deviations from the mean.

### Exposomic variables

We deliberately selected lifestyle-related variables that are known to affect some of the selected traits to construct the exposome in this study. These variables include smoking, alcohol intake, physical activity, and dietary composition. Details of these variables are listed in Supplementary Table 6. Our aim here is not to cover a comprehensive set of exposomic variables, but to demonstrate the potential use of the proposed integrative analysis of genomic and exposomic data for partitioning phenotypic variance and phenotypic prediction.

### Statistical Models

We used LMMs to simultaneously model the random effects of the genome and the exposome. The model equations and their assumed sample variance-covariance structures are summarized Table 2. In these models, a genomic relationship matrix (**G**) was constructed using an n x m_1_ genotype coefficient matrix (**A**) as 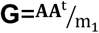, where n is the number of participants and m_1_ is the number of SNPs. Similarly, an exposomic relationship matrix (**E**) was estimated using an n x m_2_ exposomic variable matrix (**B**) as 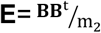 where m_2_ is the number of exposomic variables (Table 2). These relationship matrices were used to estimate the additive effects of the genome and the exposome. In addition, interaction effects, including gxe, exe and exc, were also considered in these models (Table 2).

### Principal component-based transformed variables for E

If the degree of correlation among variables is high, it can cause biased estimates when the variables are fitted in a model, i.e. multicollinearity problem. Such bias is also problematic when using correlated exposomic variables to construct **E** to be fitted in an LMM to estimate the proportion of the variance explained by the variables (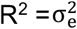 when phenotypes are standardised with mean zero and variance one). The R^2^ can also be obtained from a linear model, i.e. the coefficients of determination. For problematically correlated variables, principal component regression has been introduced^45^.

A linear model can be written as

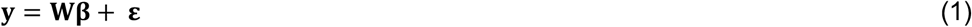

where y is a N vector of phenotypes, **W** is a column-standardised N x M matrix having correlated exposomic variables, **β** is their effects and **ε** is a vector of residuals.

When exposomic variables in **W** are highly correlated, estimated exposomic effects (**β-**hat) are inflated due to multicollinearity problem.

From the singular value decomposition, **W** can be expressed as

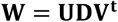

where **U** is a matrix whose columns contain the left singular vectors of **W, D** is a diagonal matrix having a vector containing the singular values of **W** and **V** is a unitary matrix (i.e. **VV**’=I) whose columns contain the right singular vectors of **W**.

V can be also obtained from the eigen decomposition of the covariance matrix of the variables, i.e. **W**^t^**W**.

The principal component regression approach^45^ proposes to transform W to a column-orthogonal matrix, Ω, multiplied by V, which can be written as

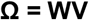

Now, we can replace W with Ω in the model as

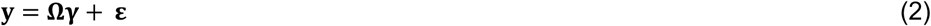

where 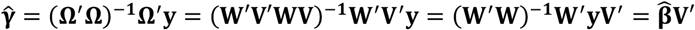

Therefore, R^2^ values obtained from models (1) and (2) are equivalent as

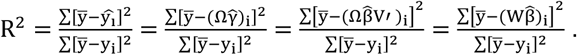

However, equation (2) can avoid a collinearity issue among the variables. Therefore, model (2) can be extended to a linear mixed model, i.e. the covariance structure can be constructed based on Ω, i.e. ΩΩ’/m where Ω is column-standardised.

Suppose a LMM of the form

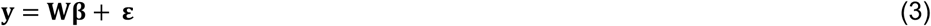

where **y** is a vector of phenotypes for n individuals; **W** is a n x m matrix that contains m exposomic variables; **β** is a vector of random exposomic effects, each assumed normal with mean zero and variance 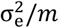 and **ε** is a vector of residuals, each assumed normal with mean zero and variance 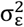

Under this model, phenotypic variance is partitioned as

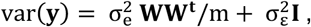

where **I** is the n x n identify matrix.

When exposomic variables are highly correlated, a transformed **W**, denoted as **Ω**, should be used, to avoid biased 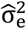.

In a similar manner to the linear models (1) and (2), LMM (3) can be rewritten as

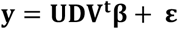

Since **V**^t^**V** = **I**

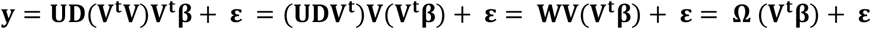

Then

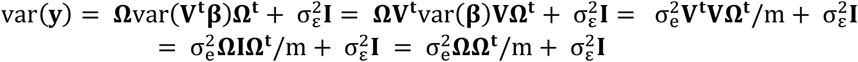

Therefore, using column-standardized principal components of exposomic variables as **W** for Equation (3) can avoid biased 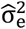. This is further verified using simulations.

### Estimation of exc interactions

We extend the proposed model to a reaction normal model (RNM) by introducing exc interaction terms, where e is the additive effects of exposomic variables and c is a covariate. Given the robust additive effects found in the above, the interest of fitting RNMs is determine whether these effects vary depending on covariates, which would be evidenced by the presence of significant exc interactions.

While estimates of 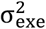 inform the phenotypic variance explained by the sum of all possible combinations of pair-wise interactions between lifestyle-exposomic variables, it may also be of interest to estimate the modulated exposomic effects specific to particular covariates, using the reaction norm model (RNM^31-33,46^). The covariates include alcohol intake, smoking, energy intake, physical activity, sex, socio-economic status (indexed by Townsend deprivation index), age and ethnicity measured using the first two ancestry principal components. For each covariate, we fitted the RNM that allows the covariate to interact with exposomic effects and compared the fit of this model with a null model that assumes no exc interactions (see Supplementary Table 3 for p-values). Significant covariates were then included in a subsequent analysis of RNM that fit multiple covariates simultaneously. We reported the total variance of exc interaction effects in Supplementary Table 7.

### Five-fold cross-validation

Using 5-fold cross validation, we 1) validate significant variance components identified above (Table 1) and 2) evaluate the extent to which the inclusion of these variance components improves phenotypic prediction. For every trait, we randomly split the sample into a discovery set (∼80%) and a target set (∼20%) and iterated this process five times in a manner such that target sets did not overlap across iterations (see Figure 6 for an outline). We derived the prediction accuracy of each model by averaging the Pearson’s correlation coefficients between the observed and predicted phenotypes across target sets; then compared prediction accuracies between models (e.g., y = g + ε vs. y = g + e + ε) to determine phenotypic prediction improvements gained by the inclusion of a given variance component (e.g., var(e)). For each variance component, we regressed prediction accuracy improvements on estimates of the variance component and declared the variance component valid if the slope differs from zero.

**Figure 6.**
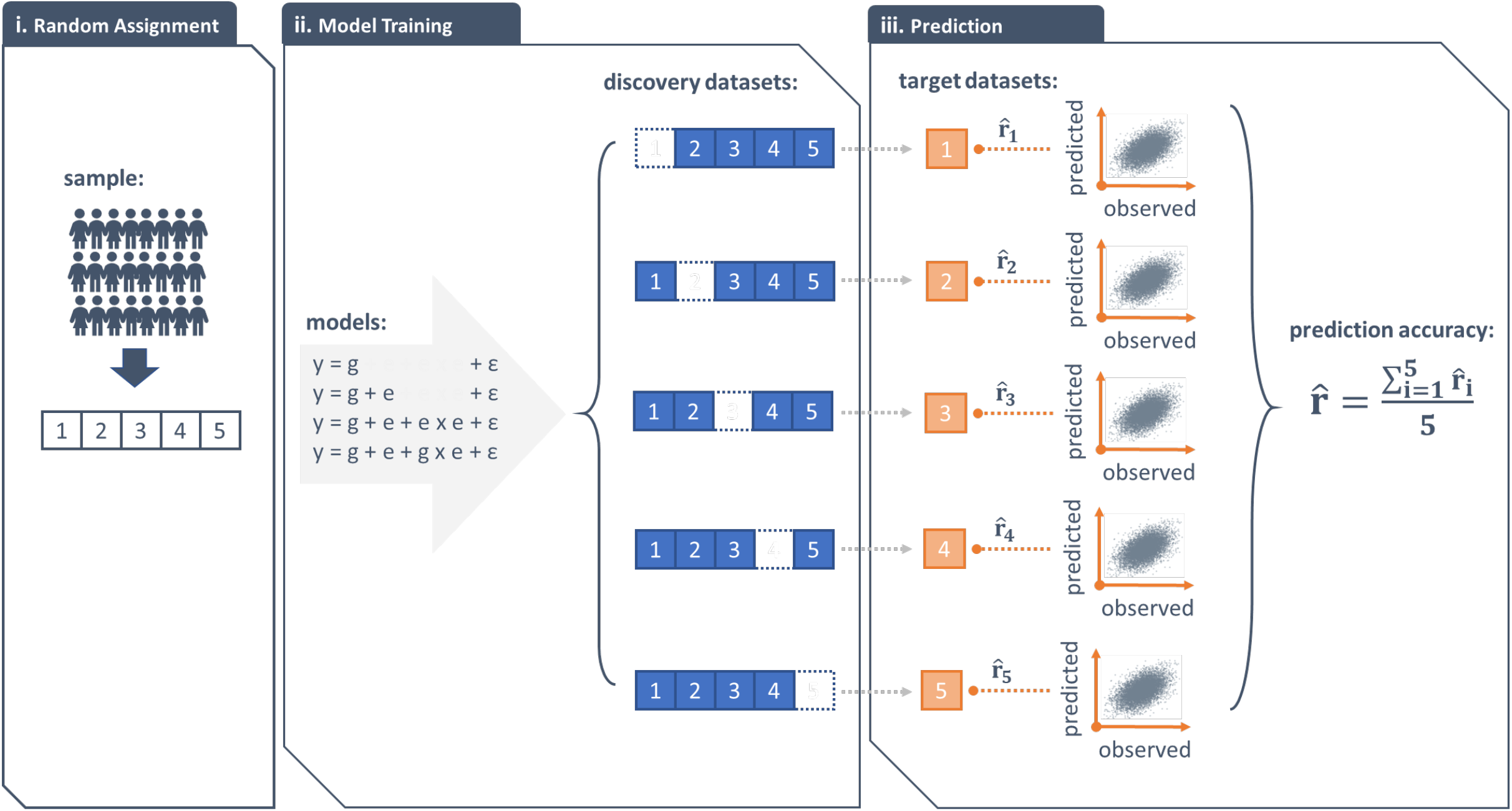
A schematic showing 5-fold cross-validation procedures. i) Randomly assign individuals to 5 groups of an equal size. ii) Choose one group as the target dataset and the remaining four as the discovery dataset. Iterate the selection process five times in such a way that target datasets do not overlap across iterations. Fit 4 models to each discovery dataset. iii) For each model, generate the best linear unbiased predictions from discovery datasets and project them onto their corresponding target datasets to derive predicted phenotypes. Compute the phenotypic prediction accuracy for each model by averaging Pearson’s correlation coefficients between the predicted and the observed phenotypes across target datasets.

### Theoretical prediction accuracy for quantitative traits

Suppose we predict phenotypes of a quantitative trait (e.g., BMI) with SNP-based heritability h^2^ using a discovery dataset of N individuals. Following previous theoretical derivations^23,27-30,34^, the genomic prediction accuracy based on the model y = g + ε can be written as

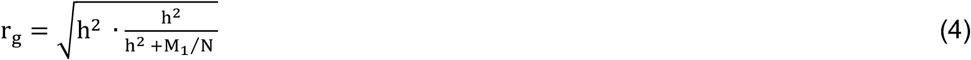

where M_1_ is the effective number of chromosome segments, which is a function of the effective number of population size and can be estimated using the inverse of the variance of genomic relationships (i.e., **G** in Table 2) between the discovery and target samples^27-30^.

Assuming that phenotypes are standardized to have mean zero and variance one, if the total amount of phenotypic variance explained by the exposome is 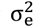 Equation 4 can be adapted to describe the prediction accuracy of the model y = e + ε in the form

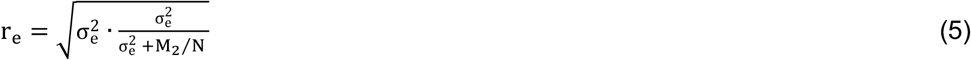

where M_2_ is analogous to M_1_ and can be thought of as the effective number of (independent) exposomic variables. Similar to M_1_, M_2_ can be estimated using the inverse of the variance of exposomic relationships (**E** in Table 2) between the discovery and target samples.

Upon establishing an agreement between expected accuracies, based on Equations 4 and 5, and observed accuracies for the 11 traits in this study (Supplementary Figure 4), we proceeded to the prediction accuracy of the proposed integrative analysis of genomic and exposomic data.

Assuming that the genomic and exposomic effects on phenotypes are uncorrelated, the prediction accuracy of the model y = g + e + ε can be written as

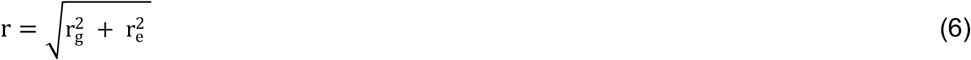

Equation 6 is verified by an agreement between the expected and observed prediction accuracies for the 11 traits in this study (Supplementary Figure 4).

### Theoretical prediction accuracy for disease traits

Considering a disease trait with a population prevalence k, we derived the expected prediction accuracy of the model y = g + e + ε for the disease in terms of the correlation coefficient between the true underlying disease liability and predicted values from the model^23,28,34,47^, which can then be converted to an AUC value^23-25^.

Similar to r_g_ and r_e_, the expected prediction accuracies for the disease on the liability scale, denoted as 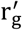 (for y = g + ε) and 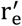 (for y = e + ε), can be computed using previous derivations^23,28,34,47^ as the followings.

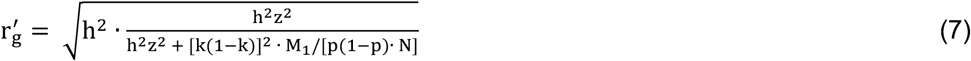

where h^2^ is the SNP-based heritability on the liability scale, N is the discovery sample size, k is the population prevalence, p is the ratio of cases in the discovery sample, and z is the density at the threshold on the standard normal distribution curve.

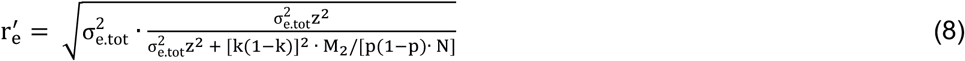

where 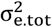 is the total amount of variance explained by the exposome on the liability scale (i.e., 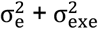. Note 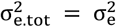 when 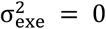.

As in Equation 6, we combined 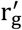 and 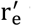 to derive the expected prediction accuracy on the liability scale for the disease, denoted as r^’^, under the assumption that the genetic effects and exposomic effects are uncorrelated.

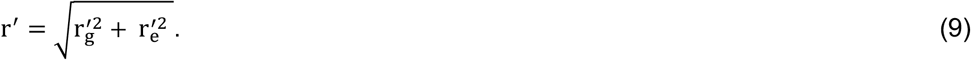

Following a well-established theory^23-25,28^ that has been verified by a comprehensive analysis of real data^26^, we converted r’ to the area under the receiver operating characteristic curve (AUC) as

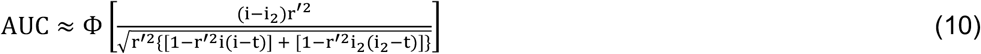

where i (=z/k) is the mean liability for diseased individuals, i_2_ (=-ik/(1-k)) is the mean liability for non-diseased individuals, t is the threshold on the normal distribution that truncates the proportion of disease prevalence k and Ф is the cumulative density function of the normal distribution.

To derive the AUC values shown in Figure 5, we set p = k, M_1_ to 50,000 and M_2_ to 28. M_1_ (50,000) was estimated from the inverse of the variance of genomic relationships (**G**) between the discovery and target samples^27,29,30^. Similarly, M_2_ (28) was estimated from the inverse of the variance of exposomic relationships (**E**) between the discovery and target samples, which agrees with the number of transformed exposomic variables by a principal component analysis in this study (see the correlated exposomic variables section in Methods). Note that setting M_2_ up to 100 would not yield expected prediction accuracies that notably differ from those from setting M_2_ = 28.

## Code availability

The source code for MTG2 v2.18 and example code along with related files for fitting IGE model can be accessed without any restrictions from https://sites.google.com/site/honglee0707/mtg2 or from https://github.com/honglee0707/IGE.

