## Supplementary information for "An integrative analysis of genomic and exposomic data for complex traits and phenotypic prediction"

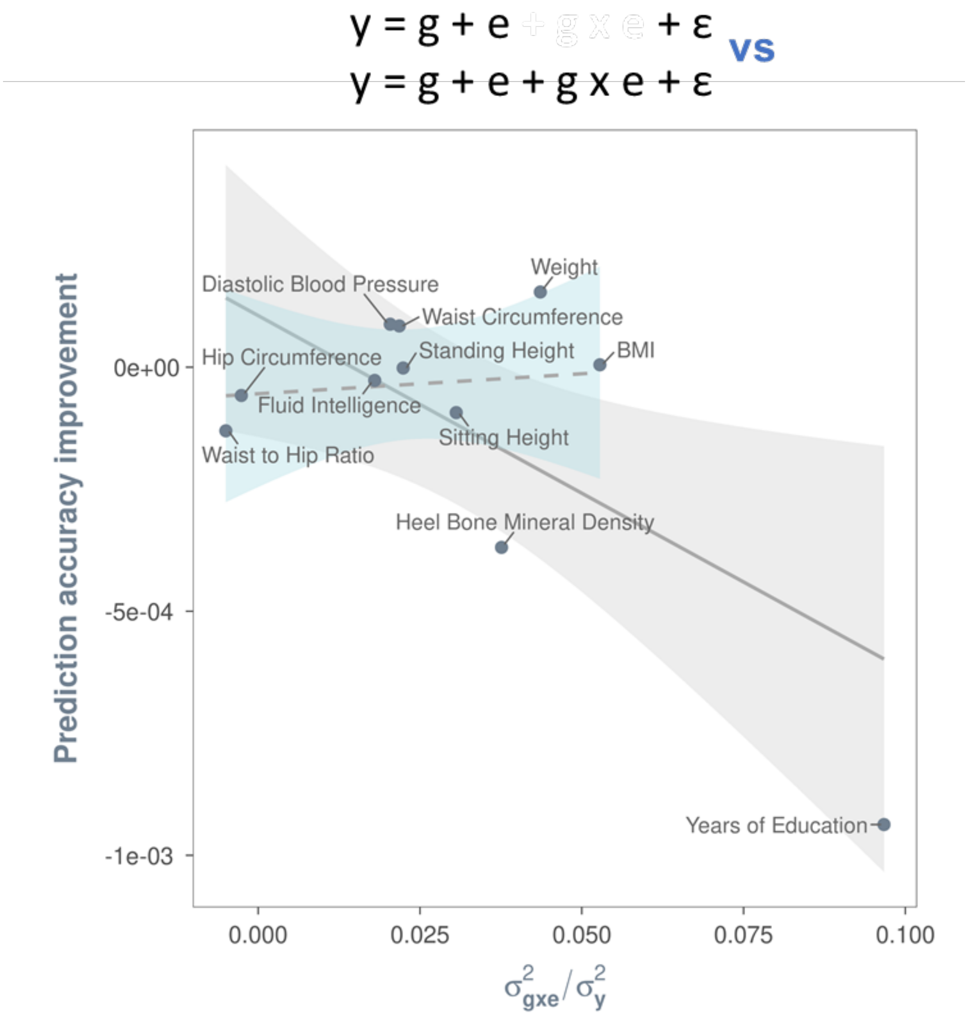


Supplementary Figure 1. Interactions between SNP genotypes and exposomic variables (gxe interactions) contribute to phenotypic variance but not phenotypic prediction accuracy. $\sigma^2_{\text{gxe}}$ denotes the phenotypic variance explained by gxe interactions. Prediction accuracy was computed using the Pearson's correlation coefficient between the observed and the predicted for models with and without a random term for phenotypic effects of gxe interaction, denoted as y = g+e+ε and y = g+e+gxe+ε, respectively. g = phenotypic effects of the genome; e = phenotypic effects of the exposome; ε = residuals. Prediction accuracy improvement (i.e., y-axis) was derived by subtracting the prediction accuracy of the model y = g+e+ε from that of the model y = g+e+gxe+ε. Least squares lines with 95% confidence band are based on a linear model that regressed prediction accuracy improvements on estimates of variance explained by gxe interactions. The solid line is based on all traits, and the dashed line is based on traits without years of education. The slope of the solid line is statically different from zero (t(7)=0.02), but the slope of the dashed line is not (t(6)=0.78).

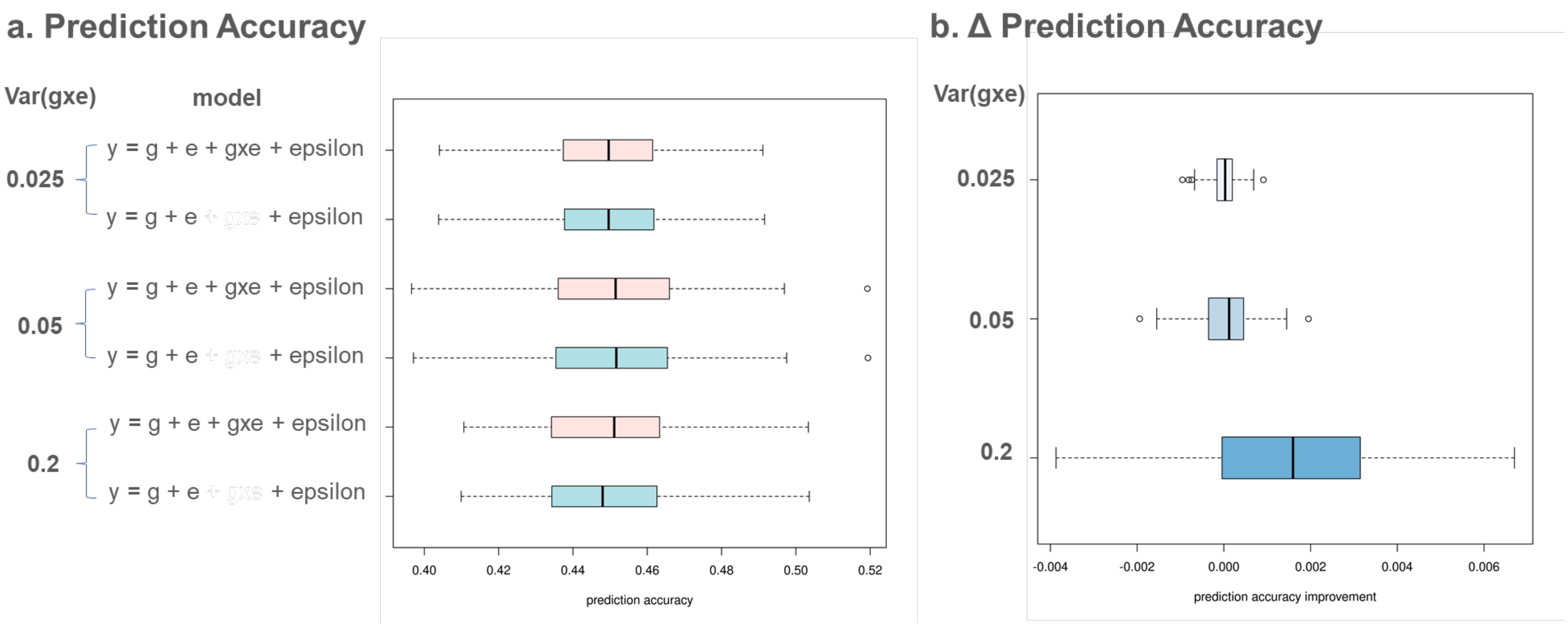


Supplementary Figure 2. Simulation results showing that despite the presence of geniune gxe interaction effects, little phenotypic prediction accuracy can be gained from acocunting for these interactions.

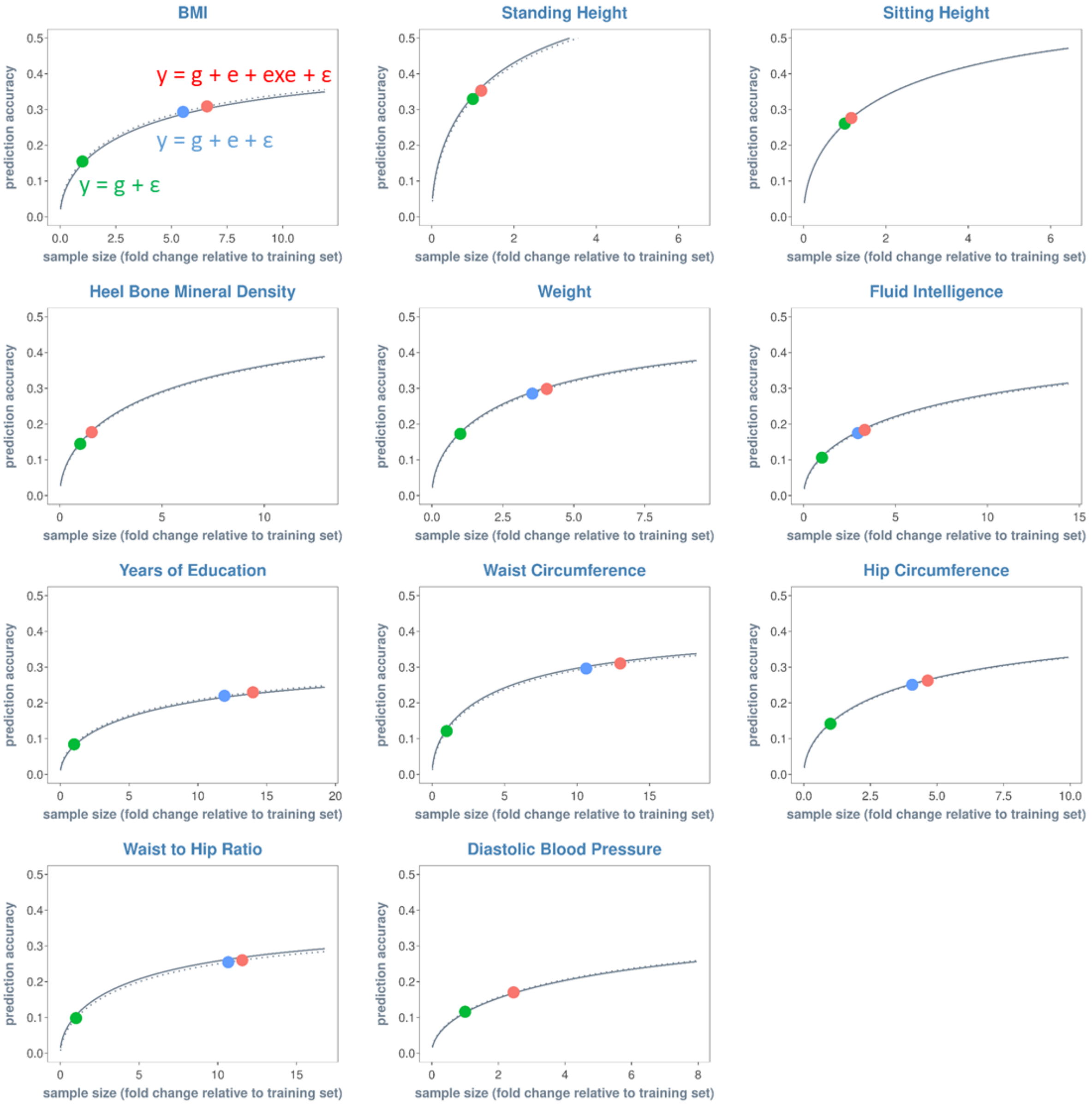


Supplementary Figure 3. Additional sample size required for the model y = g + ε to achieve the same level of prediction accuracy as y = g + e + ε (blue) and y = g + e + exe + ε (red). Sample size is expressed relative to that of the training (or discovery) dataset for each trait. Each solid curve represents the projected prediction accuracy as a function of sample size for the model y = g + ε using established theories (see Methods). Each green dot represents the observed prediction accuracy by y = g + ε at the given sample size. Dotted curves are prediction accuracies adjusted for the difference between the theoretical prediction accuracy and observed prediction accuracy, noting that the difference is minimal for all traits such that most dotted curves are obscured by the solid curve. The y-coordinates of the blue and red dots correspond to the observed prediction accuracies for models y = g + e + ε and y = g + e + exe + ε, respectively. The x-coordinates of the blue and red dots correspond to the projected sample sizes required for y = g + ε to achieve the same level of prediction accuracy as y = g + e + ε and y = g + e + exe + ε.

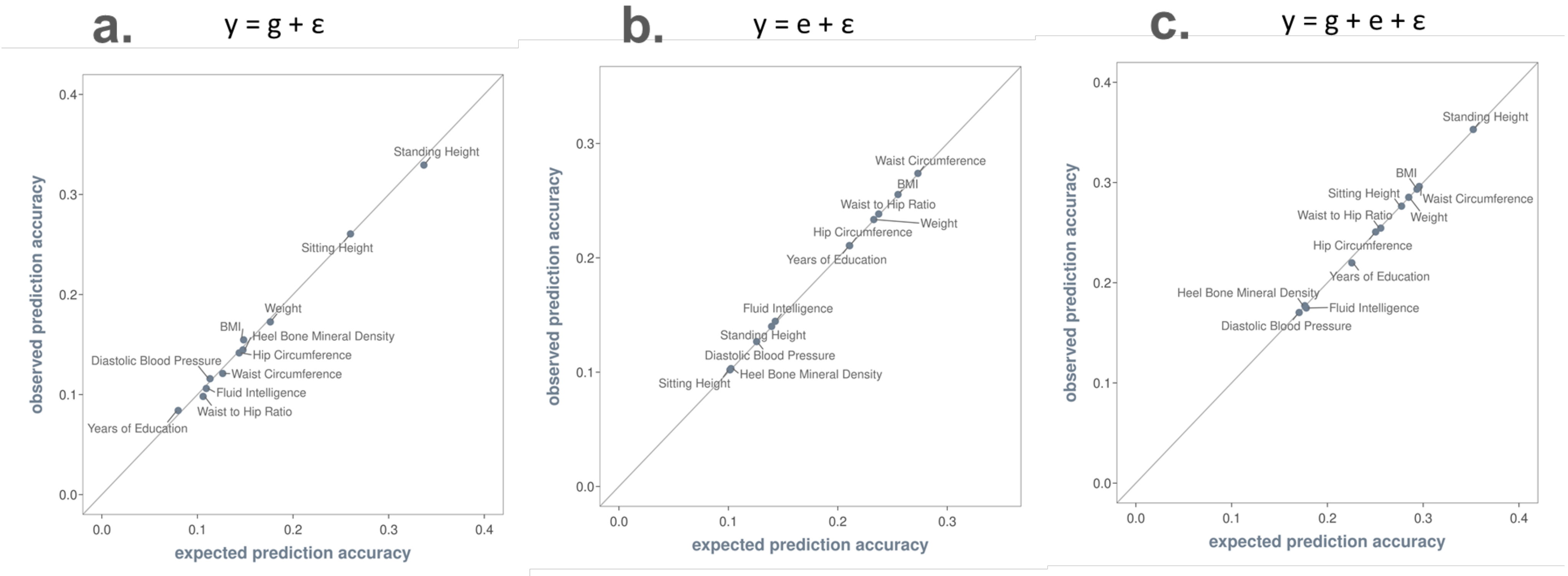


Supplementary Figure 4. Prediction accuracies derived from theories agree with prediction accuracies based on data. Expected prediction accuracies are based on established theories (see Methods). Observed prediction accuracies are based on results from 5-fold cross validations. Panel a. prediction accuracy for the model y = g + ε. Panel b. prediction accuracy for the model y = e + ε. Panel c. prediction accuracy for the model y = g + e + ε.

Supplementary Table 1. Variance and covariance estimates from model $y = g + e + \varepsilon$, where cov(g,e) is a free parameter for estimation.

| **Trait** | **var(g)** | | **var(e)** | | **cov(g,e)** | |
|---|---|---|---|---|---|---|
| | est. | s.e. | est. | s.e. | est. | s.e. |
| bmi | 1.6E-01 | 8.9E-03 | 5.2E-02 | 1.4E-02 | 2.9E-03 | 3.5E-02 |
| height | 5.0E-01 | 1.2E-02 | 1.5E-02 | 4.2E-03 | 3.5E-02 | 2.0E-02 |
| sitheight | 3.3E-01 | 1.0E-02 | 7.9E-03 | 2.3E-03 | 1.8E-02 | 1.4E-02 |
| heelbmd | 2.5E-01 | 1.8E-02 | 9.7E-03 | 3.0E-03 | -1.2E-02 | 2.0E-02 |
| weight | 2.0E-01 | 9.6E-03 | 4.4E-02 | 1.2E-02 | 2.1E-02 | 3.2E-02 |
| fluidiq | 2.0E-01 | 2.1E-02 | 2.1E-02 | 5.9E-03 | 2.9E-02 | 3.0E-02 |
| edu | 8.9E-02 | 9.8E-03 | 4.6E-02 | 1.3E-02 | -6.9E-02 | 3.2E-02 |
| waist | 1.5E-01 | 9.6E-03 | 6.6E-02 | 1.8E-02 | -1.7E-02 | 4.0E-02 |
| hip | 1.6E-01 | 8.9E-03 | 3.6E-02 | 9.8E-03 | 2.3E-02 | 2.8E-02 |
| wsthipr | 1.3E-01 | 1.0E-02 | 5.3E-02 | 1.4E-02 | -1.9E-02 | 3.4E-02 |
| diapres | 1.6E-01 | 1.1E-02 | 1.6E-02 | 4.4E-03 | -6.0E-04 | 1.9E-02 |

Supplementary Table 2. Breakdown of phenotypic variance by the model y = g + e + exe + gxe + ε.

| | $\sigma^2_g$ | | $\sigma^2_e$ | | $\sigma^2_{exe}$ | | $\sigma^2_{gxe}$ | |
|---|---|---|---|---|---|---|---|---|
| **Trait** | *est* | *se* | *est* | *se* | *est* | *se* | *est* | *se* |
| bmi | 1.93E-01 | 1.06E-02 | 7.25E-02 | 1.82E-02 | 1.88E-02 | 2.10E-03 | 4.52E-02 | 1.05E-02 |
| height | 5.18E-01 | 1.08E-02 | 1.73E-02 | 4.70E-03 | 4.00E-04 | 7.00E-04 | 2.21E-02 | 9.60E-03 |
| sitheight | 3.92E-01 | 1.10E-02 | 9.60E-03 | 2.70E-03 | 3.00E-04 | 8.00E-04 | 3.03E-02 | 1.03E-02 |
| heelbmd | 3.10E-01 | 2.17E-02 | 1.21E-02 | 3.70E-03 | 2.50E-03 | 1.90E-03 | 3.53E-02 | 1.72E-02 |
| weight | 2.37E-01 | 1.10E-02 | 5.93E-02 | 1.52E-02 | 1.60E-02 | 1.90E-03 | 3.68E-02 | 1.04E-02 |
| fluidiq | 2.05E-01 | 2.04E-02 | 2.21E-02 | 6.30E-03 | 1.14E-02 | 2.40E-03 | 1.23E-02 | 1.59E-02 |
| edu | 7.79E-02 | 9.30E-03 | 4.75E-02 | 1.24E-02 | 1.15E-02 | 1.60E-03 | 9.20E-02 | 1.16E-02 |
| waist | 1.59E-01 | 1.03E-02 | 8.47E-02 | 2.10E-02 | 1.75E-02 | 2.00E-03 | 1.35E-02 | 9.90E-03 |
| hip | 1.97E-01 | 1.06E-02 | 5.01E-02 | 1.30E-02 | 1.28E-02 | 1.70E-03 | -6.60E-03 | 9.90E-03 |
| wsthipr | 1.36E-01 | 1.01E-02 | 6.08E-02 | 1.55E-02 | 9.30E-03 | 1.40E-03 | -9.90E-03 | 9.90E-03 |
| diapres | 1.64E-01 | 1.09E-02 | 1.66E-02 | 4.60E-03 | 1.00E-03 | 9.00E-04 | 1.98E-02 | 1.11E-02 |

Note: $\sigma^2_g$ = phenotypic variance due to genetic effects; $\sigma^2_e$ = phenotypic variance due to additive effects of exposomic variables; $\sigma^2_{exe}$ = phenotypic variance due to interactions between exposomic variables, and $\sigma^2_{gxe}$ = phenotypic variance due to interactions between genotypes and exposomic variables.

Supplementary Table 3. P-values for E-C interactions estimated from single-covariate reaction normal models.

| **Trait** | age | sex | pc1 | pc2 | townsend | alc | smk2 | met_all | energy intake |
|---|---|---|---|---|---|---|---|---|---|
| bmi | 1.31E-02 | 1.34E-02 | 4.00E-01 | 8.91E-02 | 2.43E-05 | 4.33E-22 | 1.65E-32 | 5.59E-21 | 6.80E-08 |
| diapres | 3.37E-12 | 8.31E-02 | 5.76E-01 | 2.64E-01 | 1.60E-01 | 7.25E-01 | 2.56E-01 | 7.96E-02 | 4.91E-01 |
| edu | 2.67E-10 | 2.37E-02 | 3.61E-01 | 5.57E-01 | 1.12E-03 | 2.01E-06 | 1.74E-04 | 9.78E-03 | 3.70E-05 |
| fluidiq | 3.98E-01 | 7.77E-01 | 6.30E-01 | 9.42E-01 | 8.95E-02 | 2.92E-01 | 9.30E-01 | 5.62E-01 | 1.50E-05 |
| heelbmd | 1.04E-03 | 2.55E-02 | 9.53E-02 | 2.46E-01 | 9.36E-01 | 1.14E-02 | 1.78E-01 | 3.70E-03 | 1.10E-01 |
| height | 4.88E-01 | 5.38E-02 | 6.88E-01 | 5.51E-01 | 1.36E-04 | 4.88E-01 | 1.33E-01 | 1.68E-02 | 9.23E-02 |
| hip | 9.92E-03 | 1.88E-02 | 9.44E-01 | 4.34E-01 | 8.89E-04 | 5.80E-13 | 7.00E-17 | 5.74E-24 | 9.92E-06 |
| sitheight | 3.12E-01 | 3.26E-01 | 8.03E-01 | 7.80E-01 | 1.17E-03 | 5.77E-01 | 1 | 2.31E-02 | 1.37E-02 |
| waist | 1.49E-03 | 1.69E-02 | 5.35E-01 | 5.97E-01 | 2.52E-03 | 1.81E-15 | 3.87E-23 | 9.25E-37 | 3.63E-08 |
| weight | 1.24E-03 | 1.73E-02 | 2.08E-01 | 2.68E-02 | 1.56E-04 | 3.31E-16 | 2.44E-26 | 1.26E-23 | 1.89E-07 |
| wsthipr | 5.66E-02 | 1.24E-02 | 5.91E-01 | 4.85E-01 | 1.18E-01 | 1.57E-05 | 7.05E-09 | 1.70E-20 | 2.68E-05 |

Note: 1. univariate reaction norm model: $y = g + e0 + e1*c + \varepsilon$, where c = covariate (e.g., age). 2. p-values are based on likelihood ratio tests (df=1) that compare the univariate E-C interaction model with a null model ($y = g + e0 + \varepsilon$); 3. Highlighted in orange = signals remained after a familywise Bonferroni correction, where a family is defined as 9 univariate analyses for a given trait. Alpha level corrected for multiple testing = 0.05/9 = 5.56E-3.

Supplementary Table 4. The correlation structure of exposomic variables affects variance component estimates from the model $y = e + exe + \varepsilon$.

| model parameter | true value | **uncor. exp.** | | **cor.exp.** | | **pc** | |
|---|---|---|---|---|---|---|---|
| | | ave. | s.e. | ave. | s.e. | ave. | s.e. |
| v(e) | 0.4 | 0.40 | 0.004 | 0.43 | 0.012 | 0.40 | 0.003 |
| v(exe) | 0.1 | 0.10 | 0.002 | 0.10 | 0.004 | 0.10 | 0.002 |
| v(ε) | 0.5 | 0.50 | 0.001 | 0.50 | 4.5E-04 | 0.50 | 0.001 |

Note. 1. uncor. exp.: estimation based on 10 orthogonal exposomic variables simulated from a multivariate normal distribution; cor.exp.: estimation based on 10 correlated exposomic variables simulated from a multivariate normal distribution; pc: estimation based on all principal components of the 10 correlated exposomic variable. 2. The variance estimate of correlated exposomic variables (i.e., v(e) under 'cor.exp.') can be considered as the estimate from StructLMM[1]. the estimate based on transformed exposomic variables via a principal component analysis (i.e., v(e) under 'pc') can be considered as the estimate from the proposed integrative analysis of genomic and exposomic data. See supplementary note 3 for the relationship between the two models.

Supplementary Table 5. Biased estimation of the model y = e + exe + ε can be corrected by removing outliers of exposomic variables and performing a principal component analysis on exposomic variables.

| model parameter | true value | **before** | | **pc** | | **qc** | | **pc + qc** | |
|---|---|---|---|---|---|---|---|---|---|
| | | ave. | s.d. | ave. | s.d. | ave. | s.d. | ave. | s.d. |
| var(e) | 0.4 | 0.49 | 0.22 | 0.42 | 0.06 | 0.48 | 0.23 | 0.40 | 0.04 |
| var(exe) | 0.1 | 0.08 | 0.04 | 0.09 | 0.02 | 0.10 | 0.04 | 0.10 | 0.01 |
| var(ε) | 0.5 | 0.50 | 4.9E-03 | 0.50 | 0.01 | 0.50 | 0.01 | 0.50 | 0.01 |

Note. before: estimation based on ten exposomic variables from the UK biobank, which are correlated and have skewed distributions; pc: estimation based on all principal components of the exposomic variables; qc: estimation based on quality-controlled exposomic variables, of which values outside +/-3 sd from the mean are removed; pc + qc: estimation based on all principal components of the quality controlled exposomic variables.

Supplementary Table 6. Exposomic variables used to construct the kernel matrix for estimating exposomic effects on phenotypes.

| Exposomic Variable | Field ID |
|---:|:---:|
| Pack years adult smoking as proportion of life span exposed to smoking | 20162 |
| Alcohol intake (glass & pint/week) | multiple fields |
| MET minutes/week for walking | 22037 |
| MET minutes/week for moderate activity | 22038 |
| MET minutes/week for vigorous activity | 22039 |
| estimated total food weight | 100001 |
| estimated total energy intake | 100002 |
| estimated protein intake | 100003 |
| estimated total fat intake | 100004 |
| estimated carbohydrate intake | 100005 |
| estimated saturated fat intake | 100006 |
| estimated polyunsaturated fat intake | 100007 |
| estimated total sugars intake | 100008 |
| estimated dietary fibre intake | 100009 |
| estimated iron intake | 100011 |
| estimated Vitamin B6 intake | 100012 |
| estimated Vitamin B12 intake | 100013 |
| estimated folate intake | 100014 |
| estimated vitamin C intake | 100015 |
| estimated potassium intake | 100016 |
| estimated magnesium intake | 100017 |
| estimated retinol intake | 100018 |
| estimated carotene intake | 100019 |
| estimated vitamin D intake | 100021 |
| estimated alcohol intake | 100022 |
| estimated starch intake | 100023 |
| estimated calcium intake | 100024 |
| estimated Vitamin E intake | 100025 |

Supplementary Table 7. Variance estimates of exc interaction effects from multi-covariates reaction norm models.

| Trait | Covariate | var(exc) | |
|---|---|---|---|
| | | estimate | s.e. |
| bmi | townsend | 8.0E-04 | 4.0E-04 |
| bmi | alc | 9.0E-04 | 4.0E-04 |
| bmi | smk2 | 3.0E-03 | 1.0E-03 |
| bmi | met_all | 1.9E-03 | 6.0E-04 |
| bmi | energy | 1.0E-03 | 4.0E-04 |
| diapres | age | 2.0E-03 | 7.0E-04 |
| edu | age | 2.1E-03 | 7.0E-04 |
| edu | townsend | 7.0E-04 | 4.0E-04 |
| edu | alc | 7.0E-04 | 4.0E-04 |
| edu | smk2 | 5.0E-04 | 3.0E-04 |
| edu | energy | 8.0E-04 | 4.0E-04 |
| fluidiq | energy | 1.7E-03 | 8.0E-04 |
| heelbmd | age | 1.4E-03 | 7.0E-04 |
| heelbmd | met_all | 1.4E-03 | 7.0E-04 |
| height | townsend | 9.0E-04 | 4.0E-04 |
| hip | townsend | 6.0E-04 | 3.0E-04 |
| hip | alc | 1.0E-03 | 5.0E-04 |
| hip | smk2 | 1.8E-03 | 7.0E-04 |
| hip | met_all | 2.0E-03 | 7.0E-04 |
| hip | energy | 9.0E-04 | 4.0E-04 |
| sitheight | townsend | 7.0E-04 | 4.0E-04 |
| waist | age | 7.0E-04 | 4.0E-04 |
| waist | townsend | 5.0E-04 | 3.0E-04 |
| waist | alc | 1.0E-03 | 5.0E-04 |
| waist | smk2 | 2.7E-03 | 9.0E-04 |
| waist | met_all | 3.1E-03 | 1.0E-03 |
| waist | energy | 1.2E-03 | 5.0E-04 |
| weight | age | 7.0E-04 | 3.0E-04 |
| weight | townsend | 7.0E-04 | 3.0E-04 |
| weight | alc | 8.0E-04 | 4.0E-04 |
| weight | smk2 | 2.5E-03 | 9.0E-04 |
| weight | met_all | 2.1E-03 | 7.0E-04 |
| weight | energy | 1.0E-03 | 4.0E-04 |
| wsthipr | alc | 5.0E-04 | 3.0E-04 |
| wsthipr | smk2 | 1.2E-03 | 5.0E-04 |
| wsthipr | met_all | 1.9E-03 | 7.0E-04 |
| wsthipr | energy | 1.0E-03 | 5.0E-04 |

Supplementary Table 8. Simulation results based on 10 real correlated exposomic variables

| model | true | before | | pc1 | | pc2 | |
|---|---|---|---|---|---|---|---|
| parameter | value | ave | sd | ave | sd | ave | sd |
| var(g) | 0.3 | 0.28 | 0.08 | 0.27 | 0.09 | 0.28 | 0.08 |
| var(e) | 0.3 | 0.35 | 0.16 | 0.30 | 0.02 | 0.30 | 0.02 |
| var(gxe) | 0.3 | 0.27 | 0.11 | 0.30 | 0.17 | 0.27 | 0.11 |
| var(ε) | 0.1 | 0.13 | 0.11 | 0.12 | 0.15 | 0.13 | 0.11 |

Note: before: estimation based on the original exposomic variables; pc1: the erm is constructed using the principal components (PCs) of the exposomic variables, and the kernel matrix for gxe is based on the hadamard product of the grm and the erm constructed using PCs of the exposomic variables; pc2: the erm is constructed using the principal components of the exposomic variables, and the kernel matrix for gxe is based on the hadamard product of the grm and the erm constructed using original exposomic variables.

Supplementary Table 9. Simulation results based on 10 real correlated exposomic variables under a different parameter setting from.

| model parameter | true value | **before** | | **pc1** | | **pc2** | |
|---|---|---|---|---|---|---|---|
| | | ave | sd | ave | sd | ave | sd |
| var(g) | 0.3 | 0.29 | 0.06 | 0.28 | 0.06 | 0.29 | 0.06 |
| var(e) | 0.3 | 0.35 | 0.15 | 0.30 | 0.01 | 0.30 | 0.01 |
| var(gxe) | 0.1 | 0.09 | 0.07 | 0.11 | 0.11 | 0.09 | 0.07 |
| var(ε) | 0.3 | 0.31 | 0.07 | 0.29 | 0.11 | 0.31 | 0.07 |

Note: before: estimation based on the original exposomic variables; pc1: the erm is constructed using the principal components (PCs) of the exposomic variables, and the kernel matrix for gxe is based on the hadamard product of the grm and the erm constructed using PCs of the exposomic variables; pc2: the erm is constructed using the principal components of the exposomic variables, and the kernel matrix for gxe is based on the hadamard product of the grm and the erm constructed using original exposomic variables.

Supplementary Table 10. Simulation results based on 10 correlated exposomic variables that are simulated from a multivariate normal distribution.

| model | true | **before** | | **pc1** | | **pc2** | |
|---|---|---|---|---|---|---|---|
| parameter | value | ave | sd | ave | sd | ave | sd |
| var(g) | 0.3 | 0.30 | 0.05 | 0.30 | 0.05 | 0.30 | 0.05 |
| var(e) | 0.3 | 0.32 | 0.09 | 0.30 | 0.01 | 0.30 | 0.01 |
| var(gxe) | 0.3 | 0.28 | 0.06 | 0.30 | 0.05 | 0.28 | 0.06 |
| var(ε) | 0.1 | 0.12 | 0.07 | 0.10 | 0.06 | 0.12 | 0.07 |

Note: before: estimation based on the original exposomic variables; pc1: the erm is constructed using the principal components (PCs) of the exposomic variables, and the kernel matrix for gxe is based on the hadamard product of the grm and the erm constructed using PCs of the exposomic variables; pc2: the erm is constructed using the principal components of the exposomic variables, and the kernel matrix for gxe is based on the hadamard product of the grm and the erm constructed using original exposomic variables.

**Supplementary Note 1**

### Var(e) & var(exe)

Using simulations, we identified two conditions that can cause biased variance estimates of additive effects of exposomic variables and exe interactions, which are correlations between exposomic variables and skewed distributions of exposomic variables. To show the impact of the correlation structure of exposomic variables on variance estimates of exposomic effects, we simulated, for 5,000 individuals, a set of ten orthogonal exposomic variables and another set of ten correlated exposomic variables, each from a multivariate normal distribution. Based on each set of exposomic variables, we then simulated phenotypes using the model y = e + exe + ε under the parameter setting specified in Supplementary Table 4 (column 'true value'). The simulation was repeated 100 times, resulting in 100 replicates, each with phenotypes for 5,000 individuals. For each replicate, we fitted the model y = e + exe + ε and averaged variance component estimates across replicates. Results are summarised in Supplementary Table 4. When exposomic variables are orthogonal, all variance-component estimates are unbiased. By contrast, when exposomic variables are correlated, var(e) is over estimated, although the estimate of var(exe) is unbiased.

To remedy the effect of correlated exposomic variables on var(e) estimate, we used all principal components (PCs) of the correlated exposomic variables to construct the kernel matrix for estimating var(e), and used all pair-wise interaction terms of these PCs to construct the kernel matrix for estimating var(exe). Importantly, while retaining all information of the original exposomic variables, the PCs are orthogonal to each other (Jolliffe, 1982). We find that variance estimation based on the PCs of the correlated exposomic variables are unbiased (last column of Supplementary Table 4).

To show the impact of skewness of the distributions of exposomic variables on variance component estimation, we repeated the above simulations using 10 exposomic variables from the UK biobank with skewed distributions. We also noted that these exposomic variables are correlated. As shown in Supplementary Table 5, estimation based on these exposomic variables is biased for both var(e) and var(exe). Using the PCs of these exposomic variables did not completely eliminate the bias, indicating that skewness of the distributions of exposomic variables affects variance estimation independently from the correlation structure of exposomic variables.

As a remedy, we reduced the skewness by removing outliers outside 3 standard deviations from the mean. We find that after this quality control procedure the estimate of var(exe) became unbiased; but the estimate of var(e) remained biased. These results indicate that the estimation of var(e) is sensitive to the correlation structure of exposomic variables, while the estimation of var(exe) is sensitive to the skewness of the distributions of exposomic variables. When using all principal components of the skewness-corrected exposomoic variables, all variance estimates became unbiased. Taken together, to avoid biased variance estimation of exposomic effects, it is necessary to 1) conduct quality control on the exposomic variables where values outside 3 standard deviations from the mean should be removed; and 2) transform quality-controlled exposomic variables using a principal component analysis.

### var(gxe)

We also tested the effect of the correlation structure of exposomic variables on var(gxe) estimate. To do so, we simulated phenotypes based on ten correlated (but quality-controlled) exposomic variables for 5,000 individuals using the model y = g + e + gxe + ε under parameter settings specified in Supplementary Table 8. We repeated the simulation 100 times, resulting in 100 replicates. We fitted the model y = g + e + gxe + ε to each replicate and averaged variance estimates cross replicates. Results are summarised in Supplementary Table 8. All variance components are biased when the estimation is based on the corelated exposomic variables. Using PCs of the correlated variables corrected the bias for var(e) and var(gxe) (see 'pc1' in Supplementary Table 8). This observation holds for simulations under a different parameter setting (see Supplementary Table 9) and for simulations based on 10 correlated exposomic variables whose values were simulated from a multivariate normal distribution (see Supplementary Table 10).

**Supplementary Note 2**

In the main text, we report for the real data that accounting for significant gxe interactions did not lead to phenotypic prediction accuracy improvements. We hypothesized that the power of phenotypic prediction based on gxe interactions is low. Here we investigate the power of gxe-based phenotypic prediction using simulations. Spefically, we examined, for a sample of 10,000 individuals, of which 80% serves as the training set and 20% as the target set, the extent to which varying effect size of gxe interactions can improve phenotypic prediction accuracy. To do so, we simulated phenotypes using the model y = g + e + gxe + ε with $\sigma^2_{\mathrm{gxe}}$ set to 0.2, 0.05, and 0.025, respectively. Each setting has 100 replicates, and each replicate contains phenotypes of 10,000 individuals. We randomly divided each replicate into a training set (n=8,000) and a target set (n=2,000) and subsequently computed the phenotypic prediction accuracy of two estimation models, y = g + e + ε (i.e., null model) and y = g + e + gxe + ε (i.e., full model) for each replicate. Supplementary Figure 2 presents the prediction accuracies of the two models by simulation setting (2a) and changes in prediction accuracy from the null model to the full model (2b). Despite the presence of geniune gxe interactions, little prediction accuracy is gained from accounting for these interactions, and this observation holds even under the setting with the largest gxe interactions (i.e., $\sigma^2_{\mathrm{gxe}}$ = 0.2). This observation aligns with our results from real data analyses (Figure ) and indicates that the power of phenotypic predictions based on gxe interactions is low.

**Supplementary Note 3**

Here we elaborate on the equivalence of the proposed integrative analysis of genomic and exposomic data (IGE) with StructLMM.

Quoted from Moore et al.[1], StructLMM expresses the phenotypes of a complex trait, **y**, as

$$\mathbf{y} = \underbrace{\mathbf{Xb} + \mathbf{x}\beta_G}_{\mu\mathbf{I}} + \underbrace{\textstyle\sum_{l=1}^{L}(\mathbf{x} \otimes \mathbf{e}_l)\beta_l}_{\mathbf{g} \times \mathbf{e}} + \underbrace{\textstyle\sum_{l=1}^{L}\mathbf{e}_l\alpha_l}_{\mathbf{e}} + \underbrace{\Psi}_{\boldsymbol{\varepsilon}} \quad (1)$$

where $\beta_l \sim N(\mathbf{0}, \sigma_{gxe}^2/L)$, $\alpha_l \sim N(\mathbf{0}, \sigma_e^2/L)$, and $\Psi \sim N(\mathbf{0}, \sigma_\varepsilon^2\mathbf{I})$.

Here $\mathbf{Xb}$ is the fixed effects of covariates on phenotypes, $\mathbf{x}$ stores genotypes of a given SNP, $\beta_G$ is the fixed effect or persistent effect of the SNP on phenotypes, and $\mathbf{e_1}$, $\mathbf{e_2}$, … $\mathbf{e_L}$ are the set of standardized values of environments 1, 2 to L. $\beta_l$ is the random effect of SNP x environment l interaction with a variance $\sigma_{gxe}^2/L$, $\alpha_l$ is the random effect of environment l with a variance $\sigma_e^2/L$, and $\Psi$ contains residuals with a variance $\sigma_\varepsilon^2$.

The IGE equivalent terms in Equation (1) are highlighted in the annotations. Importantly, the g x e term in StructLMM is for a single SNP, but in IGE it is aggregated across all common SNPs of the genome. Under the null hypothesis, i.e., $\sigma_{gxe}^2$=0, StructLMM reduces to

$$\mathbf{y} = \underbrace{\mathbf{Xb} + \mathbf{x}\beta_G}_{\mu\mathbf{I}} + \underbrace{\textstyle\sum_{l=1}^{L}\mathbf{e}_l\alpha_l}_{\mathbf{e}} + \underbrace{\boldsymbol{\Psi}}_{\boldsymbol{\varepsilon}} \quad (2)$$

Equation (2) becomes equivalent to a simplified IGE that assumes $\sigma_g^2 = 0$ (see Equation 2a in Table 2).
